# Evidence of thermal selection from experimental evolution in the arboviral vector *Aedes albopictus*

**DOI:** 10.64898/2026.05.22.727092

**Authors:** Ayda Khorramnejad, Umberto Palatini, Daniele Da Re, Irma Lozada-Chávez, Romina Bahrami, Hugo Perdomo, Sofia Di Castri, Roberto Rosa, Helle Aronson, Chloé Lahondère, Mariangela Bonizzoni

**Affiliations:** Department of Biology and Biotechnology, University of Pavia, via Ferrata 9, 27100 Pavia, Italy; Applied Ecology Unit, Research and Innovation Centre, Fondazione Edmund Mach, 38098 San Michele all’Adige, Italy; Center Agriculture Food Environment, University of Trento, 38122 Trento, Italy; Evo-devo and Bioinformatics groups, Institute of Computer Science and Faculty of Mathematics and Computer Science, Leipzig University, 04107 Leipzig, Germany; Department of Biochemistry, Virginia Polytechnic Institute and State University, Blacksburg, VA 24061, USA

## Abstract

The extent to which phenotypic plasticity and adaptation are coupled during ectotherm thermal evolution is poorly understood. We carried out thermal experimental evolution for 20 generations in the arboviral vector *Aedes albopictus,* an efficient invasive species that has newly conquered climatically diverse regions in the past few decades. During ‘acclimation’ (one generation of evolution) we saw accelerated development and increased reproduction that traded off against survival. After only another 10 generations, we saw adaptation in the form of major changes in mosquito fitness, metabolism and gene expression, revealing the consolidation of a temperature-dependent trade-off between reproduction and longevity. These shifts demonstrate that *Ae. albopictus* can adapt at the pace of warming. When selection was relaxed, most of the thermally shifted phenotypes reverted to control levels, revealing the importance of plasticity after prolonged evolution. Furthermore, 250 warm evolution-altered genes did not return to control levels. These genes exhibited significant negative correlation between mean and variance in warm-evolved mosquitoes, but a two-fold variance reduction without mean change in relaxed-selection mosquitoes. Both signals are consistent with the action of selection operating on a polygenic trait architecture. Ecological modelling identified egg-to-adult viability as the primary driver of thermal reproductive success, highlighting juvenile stages as a crucial control target under continued warming.

## Introduction

*Aedes* spp. mosquitoes are the primary worldwide vectors of arboviruses such as dengue, Zika and chikungunya viruses, which collectively threaten half of the world’s human population^1,2^. As small short-lived ectotherms, mosquitoes are particularly susceptible to thermal anomalies. Environmental temperature (T*_a_*) non-linearly influences mosquito’s survival, reproduction, phenology, distribution, population dynamics, and competence for viral transmission^3–7^. When exposed to short-term acute thermal stress, mosquitoes rely on reversible physiological and behavioral adjustments termed ‘acclimation’^8^. These responses span molecular to organismal processes to mitigate the risk of thermal stress and associated cellular damage^5,9^, and also include behavioral buffering and range shifts to reduce exposure to damaging temperatures^5,9^. Sustained increases in T*_a_* over long periods, including those driven by global warming or possibly encountered during new invasions, promote long-lasting physiological changes, broadly termed ‘thermal adaptation’^8^. The extent and the pace at which these changes rely on phenotypic plasticity, genetic adaptation or a combination of the two remain poorly understood^10–12^.

Rapid thermal adaptation is possible in *Aedes* spp. mosquitoes due to their short generation times and high population growth rates^13^. Accumulating evidence also shows that *Aedes* spp. mosquitoes harbor vast reservoirs of genetic variants capable of contributing to rapid phenotypic and adaptive responses^14,15^. Additionally, comparative studies using individuals from latitudinal gradients report population-level differences in some behavioral (*e.g.,* thermal preference) and life-history traits, which suggest phenotypic thermal adaptations^16,17^. However, unresolved demographic histories, genetic drift, local abiotic conditions, and site-specific microbiota are confounding factors when using field-derived samples to infer thermal adaptation. Moreover, phenotypic comparisons without temporal tracking or the integration of molecular analyses do not provide complementary insights directly linking adaptations to the evolution of thermal-related traits^10–12^. Alternative strategies such as measuring phenotypic responses and genetic variability in mosquitoes after exposure to thermal variation for one or two generations have been used to investigate the potential to adapt at the pace of warming^15,18,19^. However, several simplifying assumptions are being included in the predictive models parametrized with such experimental data, including the use of a single trait underlying adaptation as well as constant phenotypic plasticity and variance over time^15,20,21^.

Experimental evolution (EE) has emerged as a powerful approach to isolate the effects of specific environmental selective pressures and identify the underlying molecular architecture of adaptive responses^22–26^. EE studies in *Drosophila* spp. detected signs of thermal adaptation within 16-20 generations of thermal selection^26,27^. Longer selection times (>60 generations) appear to improve resolution^28,29^, but the temporal scale of thermal adaptation is largely unknown and can vary across phenotypic traits^25^ and among insect species^30,31^. Furthermore, phenotypic traits involved in thermal adaptation usually have a polygenic basis^23,25,32–34^, whereby selection on many genetic loci underpins adaptive phenotypes through both complex gene networks^33–36^ and changes in gene expression variance^37,38^. Such “adaptive polygenic architecture” is not trivial, since it produces subtle allele frequency changes and/or a reduction of phenotypic variation in polygenic adaptive traits^39–43^.

Here, we applied EE under a tropical thermal regime (32°C/26°C) for three years to the arboviral vector *Aedes albopictus,* an efficient invasive species native to Southeast Asia that has become established in regions with different climates, from temperate Europe to tropical sub-Saharan Africa, over the past sixty years^44^. We quantified nearly 30 traits tracking changes in life-history, thermal behavior and metabolic states, in addition to trajectories of mosquito fitness and gene expression at defined generational intervals in replicate lines maintained under thermal selection. We further expanded all analyses to replicate lines in which thermal selection was relaxed after 13 generations of EE to clearly discriminate between non-heritable phenotypic plasticity and thermal adaptation. Collectively, our findings show the consolidation of a temperature-dependent trade-off between reproduction and lifespan occurring within 10-15 generations of EE, strongly indicating that *Ae. albopictus* can adapt at the pace of warming within a few life cycles. Ecological models calibrated on our experimental data showed that egg-to-adult viability is a stronger driver of the mosquito net reproductive rate (R₀), in comparison to female longevity and fecundity, thereby identifying juvenile stages as a priority target for vector control under continued warming. More broadly, our findings provide relevant information for the development of climate-sensitive arboviral transmission models and advance the understanding of how plastic and heritable changes interact during rapid thermal adaptation in invasive mosquito species.

## Results

### Thermal fitness of control mosquitoes remains stable during EE

We established ten replicate colonies (Fig. 1A), which were maintained under either a tropical thermal regime (“T”: 32°C for 14 h, 26°C for 10 h; “T-lines”) or standard insectary thermal conditions, as controls (“C”: 28°C for 14 h, 26°C for 10 h; “C-lines”). Both were reared at constant humidity (70%). At generation 13 (G_13_), we relaxed thermal selection for three T-lines, generating relaxed selection (R) lines that we followed for seven additional generations. Every five generations, we quantified life-history traits tracking changes in development, lifespan and reproduction in mosquitoes evolved under tropical (*i.e.,* T.G_1_, T.G_5_, T.G_10_, T.G_15_, and T.G_20_) and control (*i.e.*, C.G_1_, C.G_5_ and C.G_10_) conditions, as well as at the first and fifth generation of relaxed selection (R.G_1_ and R.G_5_) (Fig. 1B, C; Supplementary Figures 1 and 2; Supplementary Tables 1 and 2). We further assessed physiological and behavioral traits (Fig. 1D-F; Supplementary Tables 3 and 4), along with gene expression patterns in newly emerged females from C.G_1_, T.G_1_, T.G_15_, R.G_1_, and R.G_7_ mosquitoes. We observed that life-history traits of control mosquitoes remained mostly stable over generations, with changes only in the percentages of egg-to-adult viability and larval viability (Supplementary Figure 3; Supplementary Table 1). This likely reflects routine laboratory fluctuations rather than directional changes in fitness because fitness values of controls were consistent with those we had observed for over 30 generations before EE (Supplementary Figure 3; Supplementary Table 1). We compared C.G_1_ to T.G_1_ mosquitoes to identify mechanisms of thermal acclimation, while comparisons across generations and among T, C, and R lines were used to identify thermally sensitive traits and test for adaptive responses during warm evolution.

**Figure 1:**
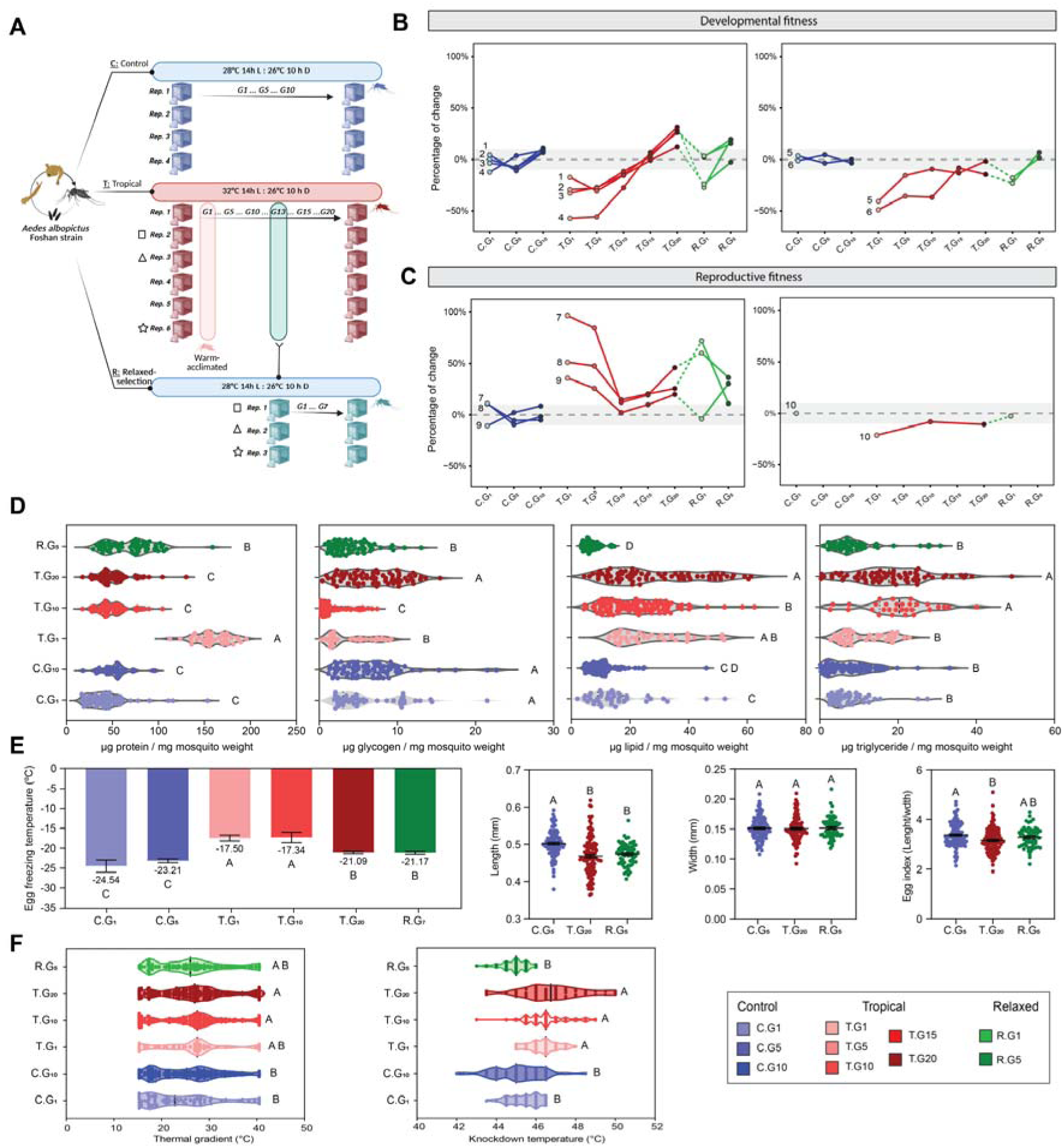
experimental design and phenotypic changes across thermal regimes and generations. (**A**) Experimental evolution design. Replicate *Ae. albopictus* lines were maintained under control (C; 28°C 14 h L : 26°C 10 h D) or tropical (T; 32°C 14 h L : 26°C 10 h D) conditions for 10 and 20 generations, respectively. At G₁₃, 1,000 eggs from three T lines were transferred to control conditions to establish relaxed-selection (R) lines, which were maintained for seven additional generations. Matching symbols (square, triangle, star) indicate the correspondence between parental T and derived R replicate lines. (**B**) Developmental fitness. Numbers label individual traits for which we show percentage changes from the mean control value. 1=female emergence time; 2=egg-to-adult viability; 3=male emergence time; 4=larval development time; 5=female lifespan; 6=male lifespan. (**C**) Reproductive fitness. Percentage change from the mean control value for female reproductive capacity. Numbers label individual traits: 7=progeny; 8=fecundity; 9=fertility; 10 = female wing length. (**D**) Energy reserves at G₁, G₁₀ and G₂₀, expressed per mg of mosquito body weight: protein, glycogen, total lipid and triglyceride. (**E**) Egg freezing temperature (mean ± SEM, values indicated below each bar) and egg morphometry (length, width, and egg index, defined as length/width). (**F**) Thermal behavior: preferred temperature along a thermal gradient (left) and knockdown temperature (KDT) (right). In B and C, dashed lines connect T lines to their derived R lines, and the grey band marks the range of variation among control lines. In D–F, letters denote compact letter display groupings; shared letters indicate no significant difference (α = 0.05). Sample sizes and statistical procedures are given in Methods and Supplementary Tables 1 to 4.

**Figure 2.**
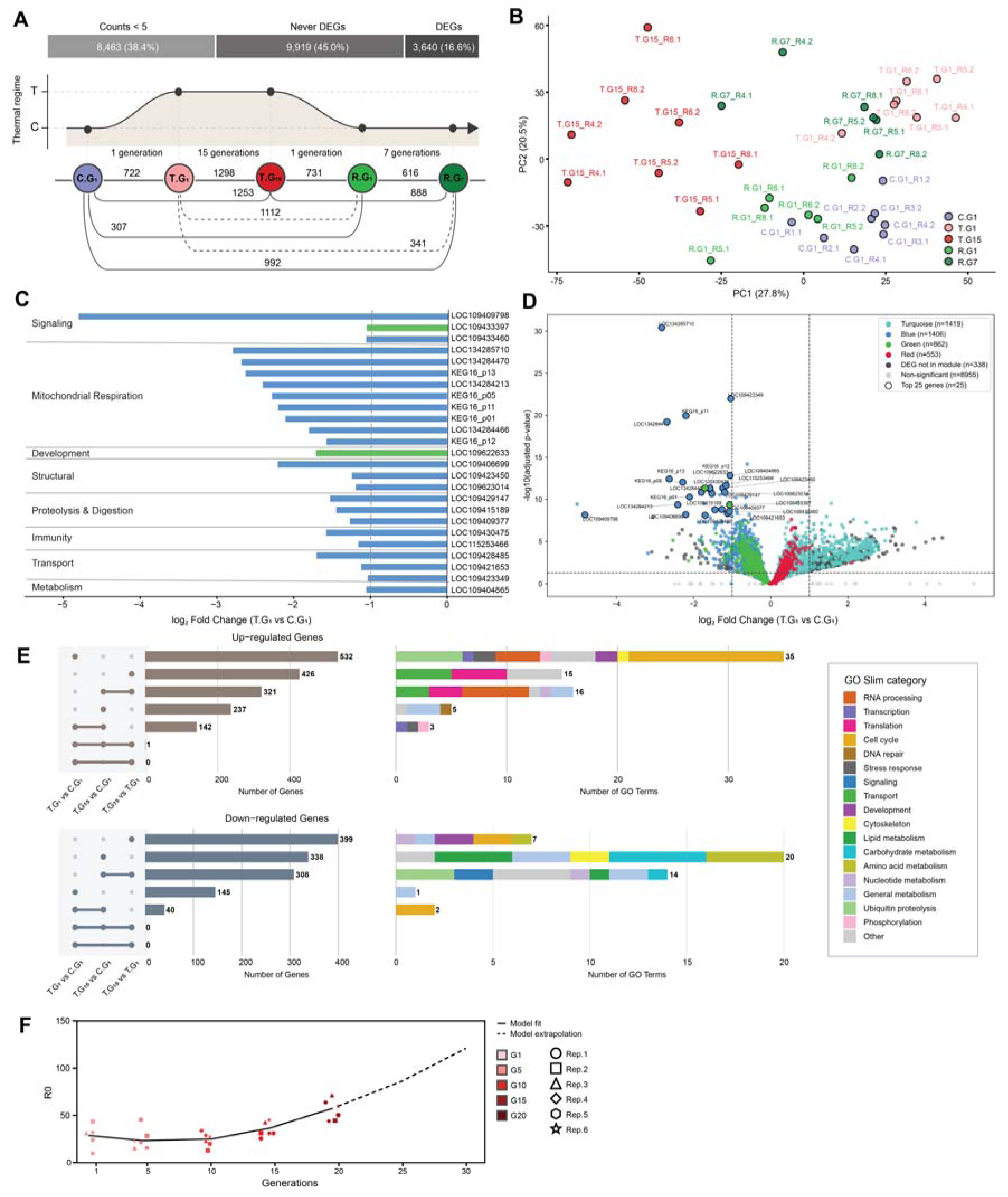
Transcriptional reprogramming during thermal acclimation and warm evolution. (**A**) Experimental timeline and differential expression across thermal conditions. Top: partitioning of the 22,022 *Ae. albopictus* annotated genes into those with mean expression count < 5 (8,463; 38.4%), those that were never differentially expressed in any pairwise comparison (9,919; 45.0%), and those that were differentially expressed in at least one pairwise comparison (DEGs; 3,640; 16.5%). Middle: thermal regimes as in Fig. 1. Bottom: number of DEGs (adjusted P < 0.01, |log₂ fold change| > 1, normalized counts ≥ 5) between control (C.G₁), warm-acclimated (T.G₁), warm-evolved (T.G₁₅), and relaxed-selection (R.G₁ and R.G₇) lines; solid and dashed lines are used to distinguish connections and do not have a specific meaning. (**B**) Principal component analysis of fat body transcriptomes (regularized log-transformed counts). Each point is one sequencing pool, labeled by replicate line. PC1 and PC2 explain 27.8% and 20.5% of the variance, respectively. (**C**) The 25 most significant DEGs between T.G₁ and C.G₁, grouped by functional category. Bar length indicates log₂ fold change; bar color indicates the assigned weighted gene co-expression network analysis (WGCNA) module. All 25 coding genes are downregulated in warm-acclimated mosquitoes. (**D**) Volcano plot of differential expression between T.G₁ and C.G₁ (log₂ fold change versus −log₁₀ adjusted P value). Genes are colored by WGCNA module assignment (turquoise, *n* = 1,419; blue, *n* = 1,406; green, *n* = 862; red, *n* = 553), with DEGs not assigned to a module in dark gray (*n* = 338) and nonsignificant genes in light gray (*n* = 8,955). The 25 genes in (C) are circled and labeled. Dashed lines mark significance thresholds. (E) Transcriptional profiles of warm-evolved (T.G₁₅) mosquitoes relative to control (C.G₁) and warm-acclimated (T.G₁) baselines. Left: UpSet plots showing intersections of upregulated (top) and downregulated (bottom) DEGs across the three pairwise comparisons. Right: stacked bars showing the number of enriched GO terms per GO Slim category for the DEGs in each intersection. (**F**) Basic reproduction number (R₀) of T mosquitoes across 20 generations of experimental evolution. Solid line: quadratic model fit; dashed line: extrapolation beyond G₂₀. Each point is one replicate line. Sample sizes, significance thresholds, and statistical procedures are given in the supplementary materials and Supplementary Tables 5, 6, 12, 13 and 14.

**Table 1.** Summary of the standardized linear model explaining variation in net reproductive rate (R₀) across generations in *Ae. albopictus*. Predictors include generation (linear and quadratic terms), female longevity, egg-to-adult female viability (EggToFemale), and fecundity. All predictors were standardized (mean = 0, SD = 1).

| Predictor | Estimate | Std. Error | t value | P value |
| --- | --- | --- | --- | --- |
| (Intercept) | 33.11 | 0.89 | 37.02 | < 0.01 |
| Generation | -9.50 | 4.30 | -2.21 | 0.038 |
| Generation<sup>2</sup> | 10.73 | 4.11 | 2.61 | 0.016 |
| Female Longevity | 6.25 | 1.35 | 4.64 | < 0.01 |
| EggToFemale | 8.92 | 1.33 | 6.72 | < 0.01 |
| Fecundity | 4.64 | 1.22 | 3.81 | < 0.01 |

### Mosquitoes maximize their reproductive capacity during thermal acclimation

We found that longevity decreased in acclimated (T.G_1_) mosquitoes compared to controls (C.G_1_), while juvenile development was faster and overall female reproductive capacity increased (Fig. 1BC; Supplementary Figure 1; Supplementary Table 2). Egg-to-adult viability showed a decreasing trend, albeit not significant. We also found that egg hatchability, larval viability, pupation rate, pupal development time (PDT), and sex ratio did not differ between T.G₁ and C.G₁ mosquitoes (Supplementary Figure 2; Supplementary Table 2).

Development under warmer-than-standard temperatures is expected to increase metabolic rate^45^, elevating energy demands during juvenile development and leading to faster depletion of carbohydrate and glycogen reserves in newly emerged females. This prediction matches what we observed in T.G_1_ females, which were smaller and had significantly higher protein and lipid content, but lower carbohydrate and glycogen content at emergence in comparison to C.G_1_ mosquitoes (Fig. 1D; Supplementary Table 3). Comparing the fat body transcriptome of T.G_1_ vs C.G_1_ females sampled at emergence, we identified a coordinated biphasic gene expression pattern that aligned with the observed phenotypic responses and involved 860 differentially expressed genes (DEGs), 722 of which had an expression count higher than 5 (Fig. 2,A to D; Supplementary Table 5). In T.G_1_ mosquitoes, we observed a general downregulation of core metabolic pathways, mitochondrial respiration, glycolysis, lipid catabolism and cellular signaling networks, including mitogen-activated protein kinase (MAPK), GTPases and vesicular trafficking, all of which are consistent with energy conservation (Supplementary Figures 4 and 5; Supplementary Tables 5 and 6). Simultaneously, genes linked to gametogenesis and egg development, as well as opsin genes, the transcription factor Glass and caldesmon, were upregulated in T.G₁ mosquitoes (Supplementary Figure 4; Supplementary Tables 5 and 6). We also observed enrichment of genes encoding germline-specific P-granule components (turquoise module in Supplementary Figure 4; Supplementary Table 6). This transcriptional profile is consistent with reduced investment in somatic maintenance and resource reallocation towards reproduction, which aligns with the trade-offs that we also observed at the fitness level with decreased longevity but increased female reproductive capacity. The observed changes in energy reserves and gene expression also support a switch from a carbohydrate-based to a lipid-based metabolism in acclimated mosquitoes. This result explains the apparent paradox of suppressed energy metabolism alongside enhanced reproductive output, because reproduction in mosquitoes relies primarily on lipid reserves (for vitellogenesis) rather than immediate ATP generation^46^.

Concurrently, we saw significant upregulation of ribosome biogenesis and the DNA replication machinery (Fig. 2, B to C; Supplementary Figure 4; 56 rRNA processing genes, P=6.4×10^-15^), which provides the molecular basis for the accelerated development seen in T.G_1_ mosquitoes. This enhanced translational capacity helps explain the reduced larval developmental time (LDT) and earlier emergence of T.G_1_ vs C.G_1_ mosquitoes (Fig. 1B). However, the shortened developmental window of T.G_1_ mosquitoes constrains tissue growth, resulting in smaller adults (Fig. 1C and Supplementary Figure 1). Furthermore, ongoing oxidative stress in T.G_1_ mosquitoes is supported by the upregulation of eight detoxification-associated genes and the coordinated suppression of nine electron transport chain subunits spanning complexes I, III, and IV, which are among the top 25 most significant DEGs (Fig. 2C; Supplementary Table 5). These transcriptomic changes, including upregulation of DNA repair pathways, are consistent with heat-induced genotoxic stress^47^ (red and turquoise modules in Supplementary Figure 4; Supplementary Table 6). Overall, these results demonstrate that *Ae. albopictus* rapidly acclimates when exposed to tropical conditions by trading lifespan for accelerated development and maximized reproduction. This coordinated response is robust across fitness, physiological and transcriptomic profiles.

### Major signs of thermal adaptation in mosquitoes emerge at 10-15 generations of EE

We monitored mosquito fitness every five generations across 20 generations of EE under a tropical thermal regime. We observed that juvenile developmental traits (*i.e.,* LDT, egg-to-adult viability, FET and MET) decreased significantly within the first five generations under tropical conditions, but they reverted to values similar to those of control mosquitoes by the tenth (T.G_10_) and/or fifteenth (T.G_15_) generation of EE (Fig. 1, B and C; Supplementary Figure 1; Supplementary Table 2). Cluster analysis resolved this as a distinct convergent trajectory (Supplementary Figure 6; Supplementary Table 7; slope = +0.132, p = 2.1×10⁻¹⁵). In contrast, female reproductive capacity showed an opposite trajectory after the thermal acclimation period: fecundity, fertility and progeny per female gradually decreased across generations under tropical conditions to reach values similar to those of control mosquitoes by T.G_10_ (Fig. 1C; Supplementary Figure 1; Supplementary Table 2) showing a consistent diverging trajectory different than the one from developmental traits (Supplementary Figure 6; slope = −0.050, p = 8.9×10⁻⁴; the two trajectories differ significantly, generation × cluster p = 3.9×10⁻⁸). We also found that phenotypic changes in females correlated with shifts in mosquito size and energy reserves. As generations under warm evolution progressed, females became larger and exhibited protein content comparable to control mosquitoes but higher lipid and triglyceride content (Fig. 1D; Supplementary Tables 2 and 3). These trends indicate that the reduced fecundity of warm-evolved mosquitoes does not reflect energetic constraints but is most likely an adaptive response to tropical conditions.

We also observed that a large proportion of mosquitoes maintained under tropical conditions died within one day of emergence up to G_10_, while a gradual increase in daily emergence time and a decrease in longevity occurred among surviving mosquitoes (Supplementary Figures 1 and 7; Supplementary Tables 8 and 9). This result suggests an initial maladaptive response to tropical conditions, which can foster adaptive divergence^48^. Adjustment of adult lifespan occurred within five generations under tropical conditions in females and then stabilized. In contrast, male longevity peaked and stabilized by the 15^th^ generation of EE, supporting the conclusion that males are less thermally resilient than females (Fig. 1B; Supplementary Figure 1). The reduction in female reproductive capacity coupled with their increased longevity during warm evolution limited the number of mosquito generations we could obtain, from eight to six generations per year, well below the 13 generations predicted for tropical climates^49^. As seen during acclimation, we also found that egg hatching time, larval viability, PDT, pupation rate and sex ratio did not change during warm evolution, revealing that these traits are not thermally sensitive under our tropical regime (Supplementary Figure 2; Supplementary Table 2).

The phenotypic responses of warm-evolved mosquitoes were mirrored by extensive transcriptional reprogramming (Fig. 2, A and E). We observed a shift from the stress response functions elicited in acclimated mosquitoes (*i.e.,* DNA repair mechanisms, detoxification, ubiquitin-mediated proteolysis) to an enhancement of translation-associated activities and a down-regulation of sugar, chitin and lipid catabolism in warm-evolved mosquitoes (Fig. 2E; Supplementary Figures 8 and 9; Supplementary Tables 10 and 11). Protein synthesis activities (*i.e.,* cytoplasmic translation initiation, small nucleolar RNAs and ribosomal small subunit assembly) were uniquely upregulated in T.G_15_ with respect to C.G_1_ (237 DEGs) or T.G_1_ (426 DEGs) mosquitoes; and they were also enriched in the 321 upregulated genes shared between the two regime comparisons (T.G_15_ vs T.G_1_ and T.G_15_ vs C.G_1_) (Fig. 2E; Supplementary Tables 10 to 12). Catabolism of sugar, chitin, and lipids, along with cell division, were enriched among the downregulated genes in T.G_15_ with respect to both C.G_1_ (338 DEGs) and T.G_1_ (399 DEGs) mosquitoes, as well as in the 308 downregulated genes shared between the two comparisons (T.G_15_ vs T.G_1_ and T.G_15_ vs C.G_1_) (Fig. 2E; Supplementary Tables 12 and 13). These results align with the observed increase in juvenile developmental time, changes in body size, and the increase in glycogen, lipid and triglyceride content at emergence of warm-evolved mosquitoes (Fig. 1, C to E). Notably, we observed more unique than shared DEGs between the compared regimes, T.G_15_ vs C.G_1_ and T.G_1_ vs C.G_1_ (Fig. 2E), which further emphasizes the difference between the transcriptomes of warm-evolved and acclimated mosquitoes. Yet, shared DEGs between both regime-sets were enriched in basal metabolic functions, including genes encoding chorion peroxidases and controlling chromosome organization (Supplementary Tables 12 and 13). Co-expression network analysis corroborated these results; modules associated with DNA damage repair were found upregulated during acclimation, but showed reduced expression in warm-evolved mosquitoes, and a module linked to transcription and translation was specifically upregulated in warm-evolved mosquitoes (Supplementary Figures 8 and 9; Supplementary Table 14).

Together, these results demonstrate that *Ae. albopictus* life-history traits gradually adjusted under continued thermal selection. Depending on the trait, values stabilized or reached slightly higher values than those of control mosquitoes. In contrast, fat body transcriptome and energy reserves shifted to a new state defined by increased lipid and triglyceride storage, enhanced biosynthesis and repressed oxidative catabolism.

### Egg-to-adult viability drives mosquito reproductive success during warm evolution

We calculated the net female reproductive rate^50^ (R₀; see equation 1, the average number of female offspring produced by a parental female) across all T replicate lines and every five generations to quantify how the reproductive success of mosquitoes changed during warm evolution (Fig. 2F). We observed that while R₀ initially dropped through G_5_, it recovered by G_15_ (Fig. 2F). Modelling R₀ across generations using a quadratic effect (see equation 5; adjusted R² = 0.55, F(2,26) = 18.02, p < 0.001) revealed a marginally negative, but nonsignificant linear effect of generations (β = –2.18, p = 0.054) and a significant positive quadratic effect (β = 0.166, p = 0.002). The vertex of the fitted parabola occurred at approximately generation 6.6, suggesting that R₀ initially declined during warm evolution as part of an early maladaptive response to tropical conditions, before increasing again after G_6_ (Fig. 2F; Supplementary Table 15). R₀ captures composite reproductive success and emerges from the interplay of multiple components, all of which may respond differently to temperature. To identify which traits most strongly contribute to variation in R₀ across generations, we fitted a multivariate linear model with linear terms for female longevity, fecundity, and egg-to-adult viability as a proxy of juvenile development, and a quadratic term for the generation (see equation 6; Table 1). Among the tested variables, egg-to-adult viability emerged as the strongest driver of R₀ across generations (standardized β = 8.92, p < 0.001). In contrast, the standardized effect sizes of fecundity and longevity were comparatively smaller (β = 4.64 and 6.25, respectively).

### Disentangling plastic and adaptive responses during thermal adaptation

To distinguish if phenotypic changes in warm-evolved mosquitoes are most likely the outcome of recurrent, non-heritable, plasticity or the emergence of evolutionary adaptations, we relaxed thermal selection and quantified fitness traits and transcriptome changes in mosquitoes under relaxed selection at both G_1_ and G_5-7_. This approach allowed us to identify a potential acclimation effect during the first relaxed generation (R.G_1_) back to control conditions. In R.G_1_ mosquitoes, we observed shifts in juvenile development and female reproductive capacity analogous to those seen at T.G_1_, albeit less pronounced (Fig. 1, B and C). We also identified 83 DEGs shared between the two acclimation events: the initial exposure to tropical conditions (T.G_1_ vs C.G_1_) and the return to control conditions (R.G_1_ vs T.G_15_) (Supplementary Figure 10; Supplementary Table 16). Of these, 63 DEGs had opposite directions of expression in T.G_1_ or R.G_1_ vs C.G_1_ mosquitoes, while showing stable expression levels across T.G_15_ lines, except for LOC134284470 and LOC134288418. In contrast, 20 DEGs showed a similar expression profile in both acclimated groups, opposite to that of T.G_15_ mosquitoes. While most of these 83 genes are uncharacterized, several have detoxification functions, including glutathione and chorion peroxidases (LOC109401758, LOC109429357, LOC109399009) and the cytochrome P450 4C1 (LOC109430500), whose role in thermal stress has been proven in *Bemisia tabaci*^51^ (Supplementary Figure 10; Supplementary Tables 16 to 18). We also identified 726 genes that were exclusively differentially expressed in R.G_1_ vs T.G_15_ mosquitoes (Supplementary Figure 10; Supplementary Table 17). Upregulated genes were enriched in activities associated with cell cycle control, while downregulated genes had mostly metabolic functions (Supplementary Figure 10; Supplementary Tables 16 to 18). Both trends suggest a decelerating pace of biochemical reactions in mosquitoes returning to control conditions, as the metabolic theory of ecology hypothesizes^4,45^.

After five generations under relaxed selection, several thermally sensitive traits (*i.e.,* egg hatchability, LDT, FET, MET, egg-to-adult viability, fecundity and progeny per female) and energy reserve contents (lipids and triglycerides) showed values not significantly different from those of both warm-evolved and control mosquitoes (Fig. 1, B to E; Supplementary Tables 2 and 3). In contrast, the global transcriptome of R.G*_7_* mosquitoes occupied an intermediate position between that of C.G_1_ and T.G_15_ mosquitoes (Fig. 2B), as measured by Euclidean distance from the control centroid (Fig. 3A). In immune-related genes, reversion to the expression levels of control mosquitoes was observed already at the first generation upon relaxing thermal selection (Supplementary Figure 11, Supplementary Table 19). Interestingly, the expression of immune-related genes was displaced downwards relative to all other genes during acclimation and warm-evolution (T.G1 vs C.G1, median Wald statistic −0.35 versus +0.07 for background, padj = 2.1 × 10^-9^; T.G15 vs C.G1, −0.46 versus +0.15, padj = 2.4 × 10^-8^) and move upwards upon returning to control conditions (R.G1 vs T.G15, +0.58 versus -0.20, padj = 1.1 × 10^-14^). These results of a thermal-dependent shift in baseline immune investment offer a transcriptional correlation of the change from resistance to tolerance observed in control vs warm-evolved mosquitoes challenged with cell fusing agent virus^6^.

**Figure 3.**
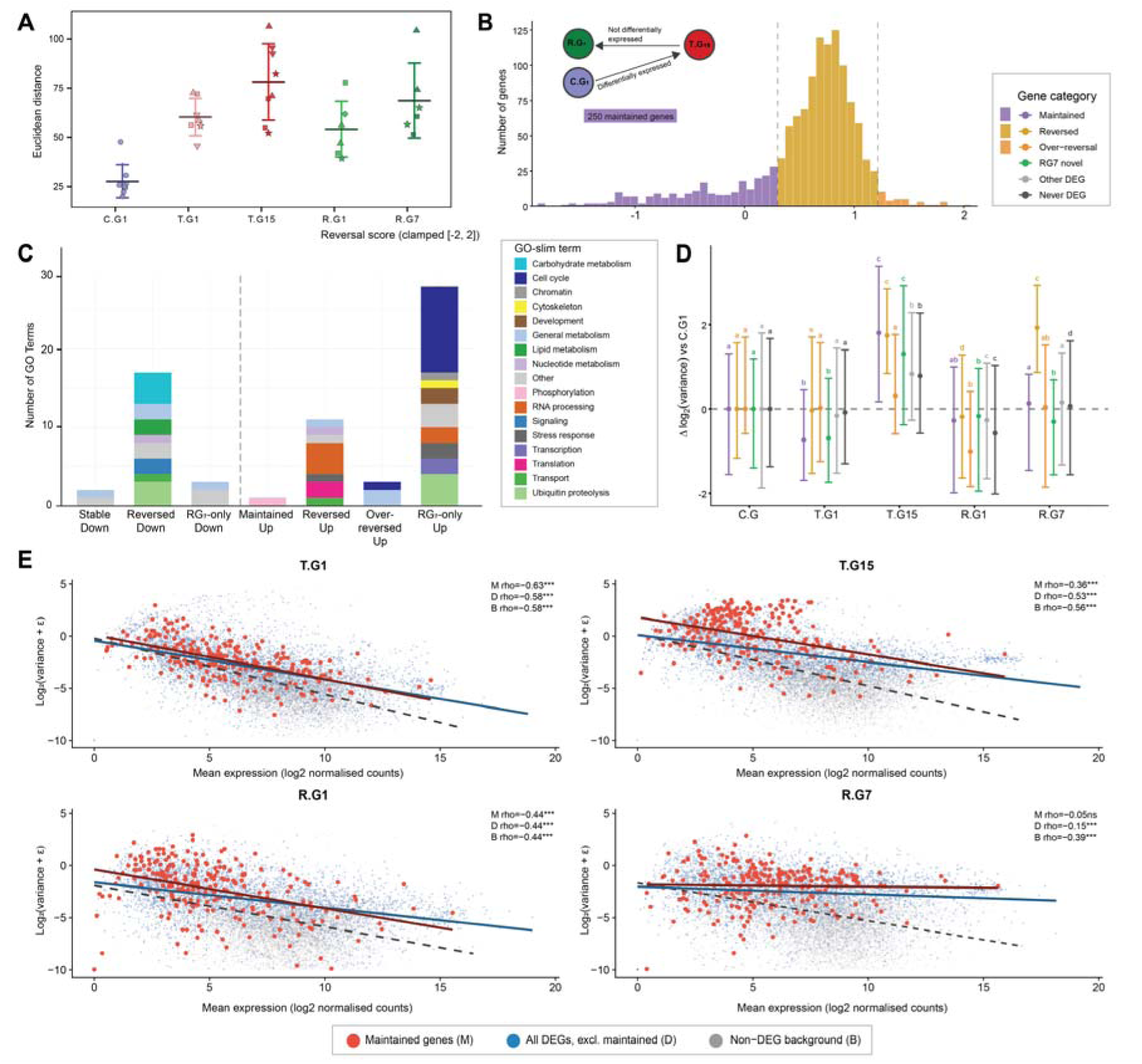
Transcriptomic reversal after relaxed selection. (**A**) Transcriptomic divergence from the control baseline, computed as the Euclidean distance of each replicate line’s fat body expression profile from the C.G₁ centroid (log₂-transformed normalized counts, genes with mean count ≥ 5). Points are individual replicate lines; symbols identify replicates; bars show group means ± SD. Colors denote experimental group: control (C.G₁, blue), warm-acclimated (T.G₁, pink), warm-evolved (T.G₁₅, red), and relaxed selection (R.G₁, light green; R.G₇, dark green). (**B**) Distribution of per-gene reversal scores for the DEGs identified in T.G₁₅ versus C.G₁. A reversal score value of 1 indicates complete reversal, 0 indicates full maintenance of the warm-evolved state, and values > 1 indicate over-reversal. Dashed lines mark the classification thresholds: maintained (purple, < 0.3; *n* = 250), reversed (yellow, 0.3 to 1.2), and over-reversed (orange, > 1.2). Inset: genes maintained in R.G₇ remain differentially expressed relative to C.G₁ and had the same expression levels as in T.G₁₅ mosquitoes. (**C**) GO Slim enrichment of DEGs stratified by reversal category and direction of regulation. Stacked bars give the number of enriched GO terms per functional category; a dashed vertical line separates downregulated (left) from upregulated (right) gene sets. (**D**) Change in inter-replicate expression variance relative to C.G₁ [Δlog₂(variance)] for each gene category across conditions. Points are category medians, bars the interquartile range. Letters denote compact letter display groupings from pairwise Wilcoxon signed-rank tests with Benjamini-Hochberg correction; within a category, shared letters indicate no significant difference across conditions (α = 0.05). (**E**) Relationship between mean expression and inter-replicate expression variance for maintained genes (M, red; *n* = 250), all other DEGs (D, blue), and the non-DEG background (B, gray) in T.G₁, T.G₁₅, R.G₁, and R.G₇ lines. Lines are linear fits per category. Spearman’s ρ is given for each category; \*\*\**P* < 0.001; ns, not significant. Sample sizes, significance thresholds, and statistical procedures are given in the supplementary materials and Supplementary Tables 20 to 22.

To better characterize the extent of transcriptional reversion in R.G*_7_* mosquitoes, we computed a per-gene reversal score for each DEG between warm-evolved and control mosquitoes (T.G_15_ vs C.G_1_, with mean normalized counts ≥10). The score quantifies how far expression in R.G₇ returns toward C.G_1_ levels relative to T.G_15_: a score of 1 indicates complete reversal, 0 indicates full maintenance of the warm-evolved state, and values >1 indicate over-reversal beyond control levels (see Methods; Supplementary Table 20). We classified 961 genes with scores between 0.3 and 1.2 as “reversed,” 250 genes with scores <0.3 as “maintained,” and 42 genes with scores >1.2 as “over-reversed” (Fig. 3B; Supplementary Table 20). We found that reversed genes upregulated in T.G15 were enriched in RNA processing, while reversed genes downregulated in T.G15 were enriched in chitin and lipid metabolism (Supplementary Table 21).

We also identified 754 unique DEGs in R.G_7_ vs C.G_1_ mosquitoes and labelled them as “R.G_7_-only” (Supplementary Table 22). Upregulated DEGs in “R.G_7_-only” were enriched in activities linked to cell cycle control, development and ubiquitin proteolysis, while the downregulated DEGs were enriched in general metabolism (Fig. 3C; Supplementary Tables 21 and 22). These results align with the enrichment in some gene ontology categories that we observed among the 726 DEGs exclusively found in R.G1 vs T.G_15_ mosquitoes, suggesting slow metabolic adjustment back to control conditions. Gradual physiological adjustment is also supported by the observation that the protein and glycogen content of R.G_5_ mosquitoes was comparable to that of T.G_1_ mosquitoes, but significantly different from that of both control and warm-evolved mosquitoes (Fig. 1D). The 250 “maintained” genes were enriched in phosphorylation activity and included 39 long noncoding RNAs (Fig. 3C; Supplementary Tables 20 and 21). We further found that the mean and variance expression values among the 250 maintained genes were significantly and negatively correlated in T.G_15_ mosquitoes, showing a two-fold increase in gene expression variance in comparison to the other thermal regimes (Fig. 3, D and E). However, such correlation was found weaker and not significant in R.G_7_ mosquitoes, accompanied by a considerable reduction of gene expression variance in comparison to the other thermal regimes (Fig. 3E).

### Egg-freezing point reached a new optimum during warm evolution

Knockdown temperature (KDT) and thermal preference (T*_p_*) assays were used to examine changes in behavioral thermal responses across generations under tropical conditions. Warm-acclimated and warm-evolved mosquitoes showed significantly higher KDT than controls, while KDT in R-mosquitoes reverted to a value not significantly different from that of control mosquitoes (Fig. 1F; Supplementary Table 4). Mosquitoes maintained under tropical conditions for at least 10 generations also showed significantly higher T*_p_* values than control mosquitoes. In contrast, warm-acclimated and relaxed-evolved mosquitoes showed T*_p_* values not significantly different from those of either C.G_1-5_ or T.G_10-20_ mosquitoes (Fig. 1F). Furthermore, control mosquitoes had the lowest egg freezing point, while acclimated mosquitoes (T.G_1_) and T.G_10_ mosquitoes showed the highest values. R.G_5_ mosquitoes had intermediate values that were not significantly different from those of T.G_20_ mosquitoes (Fig. 1E). These results align with egg morphometrics: egg length and egg index were significantly different between C.G_5_ and T.G_20_, but remained unaltered between T.G20 and R.G5 (Fig. 1E).

## Discussion

While the difference between thermal acclimation and thermal adaptation has long been appreciated for endotherms^52^, the detection of thermal adaptation in ectotherms such as mosquitoes has mainly been inferred through fitness comparisons of mosquito populations sampled at different latitudinal gradients or exposed to a thermal challenge for one or two generations^15–17^. Here, we relied on a multi-generational EE approach applied to *Ae. albopictus* mosquitoes to identify which life-history traits are thermally sensitive and investigate the interplay between non-heritable plasticity and potential adaptive changes over 20 generations of thermal selection. We investigated the thermal evolution of *Ae. albopictus* at this scale for the first time, because this species has become an increasingly relevant arboviral vector across varying climatic and ecological settings over the past century^44,53^. Diverse studies have shown that *Ae. albopictus* natural populations developed several strategies to cope with a wide range of T*_a_*^5^ and appear to have adapted within 10-30 years of population expansions^54–57^. Additionally, we recently showed that prolonged exposure to heat enhances *Ae. albopictus* tolerance to viral infection^6^. Thus, understanding the evolutionary strategies that allow this species, along with congeneric arboviral vectors, to cope with accelerated climate changes crucial to improving the forecasting of arboviral transmission risk and informing the best-suited control strategies as the climate crisis intensifies^7,20,25^.

### Robust acclimation to a new thermal condition

We found that *Ae. albopictus* mosquitoes traded longevity for accelerated development and increased reproductive output in the first generation under tropical conditions, revealing fast responses in major life-history traits during thermal acclimation. The observed changes in energy reserves and transcriptomic profile support a switch from a carbohydrate-based to a lipid-based metabolism in acclimated mosquitoes to maximize reproduction, as reproduction relies primarily on lipid reserves for vitellogenesis rather than on immediate ATP generation in mosquitoes^46^. In acclimated T.G_1_ lines, we further observed an overall enhancement of DNA repair mechanisms and detoxification genes, as well as a downregulation of respiratory chain complexes. These transcriptional changes support a metabolic shift from an efficient oxidative metabolism to a state of stress, consistent with physiological short-term responses to heat stress described across vertebrates and other invertebrates^58–62^. The observed changes in fitness trajectories during acclimation can be primarily ascribed to regulation of gene expression and non-heritable plasticity because in the first generation of EE there are no shifts in allele frequencies other than by drift, which was minimized by maintaining lines at a large population size^63^. Having used the *Ae. albopictus* Foshan strain for EE, we can also expect an absence of pre-existing adaptations to thermal stress because this strain is from the native species range in Southern China and has been maintained under laboratory conditions for over 40 years, without losing polymorphism^36^.

### A fitness trade-off between reproduction and lifespan drives thermal adaptation

We show the consolidation of a temperature-dependent trade-off between reproduction and lifespan in *Ae. albopictus* mosquitoes by the 10-15^th^ generations of EE under tropical conditions. Within the first five generations of thermal selection, we saw that a short lifespan is rapidly traded off against an increased thermal performance in juvenile development and adult reproduction (Fig. 1, B and C; Supplementary Figure 6; Supplementary Table 7). Decline in longevity was lowest in T.G_1_ in both sexes, and values increased to stabilize at control values by T.G_5_ in females and T.G_15_ in males. In parallel, female reproductive capacity (*i.e.*, fecundity, fertility and progeny per female) gradually decreased up to T.G_10_, when it started to increase again and reached slightly higher values than those of control mosquitoes by T.G_20_.

In contrast to the transcriptional stress responses elicited in thermal acclimation, fitness traits of warm-evolved mosquitoes at T.G_10_ and T.G_15_ were mirrored by extensive transcriptional reprogramming of the fat body, which converged to a metabolic state characterized by higher lipids, glycogen and triglycerides, enhanced biosynthesis, and reduced investment in cuticle remodeling and oxidative catabolism. The high mortality within one day of emergence during the first 10 generations of EE, and its rapid decline afterward, reflects strong thermal selection and support of our findings. Also, the fitness traits of control mosquitoes were stable for over 30 generations before EE, suggesting that basal trade-offs among juvenile development, adult lifespan and blood-dependent female reproduction were largely stable under our control conditions. The strong phenotypic stability of controls also argues against large-scale genetic drift, although we cannot exclude some transcriptional drift.

We suggest that such a trade-off was triggered in *Ae. albopictus* by negative fitness associations of covarying life-history traits and antagonistic pleiotropic effects on alleles affecting the allocation of resources to development or reproduction under a long-term thermal challenge. Our hypothesis aligns with findings reported in *D. melanogaster*^64–66^ and with the modeling predictions of Pawar *et al.*^67^, showing that trait trade-offs can constrain thermal adaptation in arthropods due to covariance among four key life-history traits: juvenile development, adult fecundity, juvenile and adult mortality. We thus conclude that the consolidation of a trade-off switch between reproduction and lifespan, after an acclimation period, is a strong indicator of *Ae. albopictus* being able to adapt at the pace of increased warming within a few life cycles. To the best of our knowledge, this is the first study that thoroughly demonstrates a temperature-dependent trade-off between reproduction and lifespan in invasive mosquitoes through experimental evolution.

### Plastic phenotypic responses facilitate rapid thermal adaptation

Collectively, our findings robustly show that plastic phenotypic responses can facilitate adaptive evolution to a prolonged thermal challenge in *Ae. albopictus*. Coordinated plastic changes in gene expression and life-history traits, together with metabolic rewiring, actively sustain rapid generational turnover that enhances reproductive fitness under an accelerated life cycle and allows warm-evolved female mosquitoes to return to the phenotypic state of control mosquitoes once thermal selection is relaxed. Several lines of evidence support this conclusion.

First, fitness values of several thermally sensitive developmental traits (*i.e.*, MET, FET, adult longevity, egg freezing point) remained at warm-evolved values in R.G_7_ mosquitoes, supporting the hypothesis that these traits reached new optima through the regulation of genes responsive to thermal selection. In contrast, fitness values of thermally sensitive traits for both development and reproduction (*i.e.,* LDT, egg hatchability, egg-to-adult viability, fecundity and fertility) and energy reserve contents (lipids, glycogen and triglycerides) changed in R.G_1_ mosquitoes and reversed to values of control lines by R.G_5_, suggesting that plastic changes redirected the transcriptomes to the original state of control mosquitoes during relaxed warming. Accordingly, approximately half of the gene expression profiles of R.G_7_ mosquitoes reversed to control expression values, with substantial reversion already present in R.G_1_. Since transcriptome profiling was performed after fifteen generations of EE under tropical conditions and seven generations of relaxed selection, variance in gene expression most accurately reflects fitness changes at time points where warm-evolved and relaxed-evolved states were fully induced. The fitness and transcriptomic convergence of several key mosquito traits in warm-evolved and relaxed selection lines strongly indicates that phenotypic plasticity is not affected by long-term thermal evolution, in agreement with similar findings in *Drosophila* spp.^27,68^. The role of phenotypic plasticity in facilitating the return of long-term evolved traits to basal conditions after relaxed selection was also reported in vertebrates and *Escherichia coli* for distinct long-term environmental stresses^69,70^.

Second, we found a distinct association in changes of mean and variance expression for 250 DEGs in T.G_15_ that maintained ≥ 70% of their fold-change in R.G_7_ mosquitoes, suggesting a clear differential action of natural selection in both thermal regimes. In T.G_15_ mosquitoes, we found a significant and negative correlation between the mean and variance in the expression of the 250 maintained genes, which indicates a clear signal of directional selection over phenotypic variance during prolonged thermal challenge^37,38^. This correlation is even stronger than that observed for the remaining DEGs in the same thermal regime, and also all DEGs exhibited a two-fold increase in gene expression variance relative to the other regimes (Fig. 3E). In R.G_7_ mosquitoes instead, we observed an absence of correlation between mean and variance in the expression of the 250 maintained genes, and in all remaining DEGs, accompanied by a pronounced reduction in gene expression variance (Fig. 3E). Both trends in R.G_7_ mosquitoes are most likely indicating the presence of canalization of mosquito traits due to stabilizing selection and/or directional selection operating on polygenic traits^37,38,71^. These findings are consistent with the ‘omnigenic model’ of complex traits in which peripheral genes, connected through regulatory networks, contribute collectively with core genes of major effects to complex phenotypes^32,34^. Under this scenario, core genes directly linked to thermally responsive traits are expected to contribute only a small fraction of total trait heritability, as selection acts at higher orders in polygenic phenotypes^32^.

Our findings are also in good agreement with studies reporting abundant phenotypic plasticity and genomic variation available in mosquitoes and *Drosophila* spp. to cope with thermal changes at different timescales^15,16,27,33,37,69,72–74^, which may result in similar or repeated phenotypic and gene expression changes across mosquito species^32,34,75,76^. It remains unclear whether the observed phenotypic and genetic changes contribute to organismal adaptation or are consequences of it^76^. By defining a core set of thermally sensitive phenotypes and genes (Fig. 1, B and C; Supplementary Figures 1 and 6; Supplementary Tables 2, 17, 18 and 20), this study provides specific targets on which genetically based evolutionary change can be further tested.

### Broad ecological implications

Our findings unveil the potential role of phenotypic plasticity in facilitating the adaptation of fundamental life-history traits of invasive mosquitoes during short- and long-term thermal challenges, as observed in invasive plants^77^ and other insects^78,79^ as well. This should be considered when modelling transmission risk, since phenotypic plasticity can be incorporated into models through functions describing a faster or delayed response to changes in ambient temperature^80^. Our work also highlights the challenges in detecting clear signals of thermal adaptation and thermally dependent phenotypic changes due to polygenic and covarying life-history traits and the persistence of phenotypic plasticity during warm evolution. Failing to account for these factors can confound the separation of phenotypic plasticity from thermal adaptation. This is because the phenotypic and genetic differences observed in comparisons across geographic populations or during EE without a temporal gradient can be the outcome of confounding factors, including adaptations to local abiotic conditions, acclimation, genetic drift or population structure^20,21,37,38,43,81^.

Modelling R₀ across generations revealed a trait hierarchy influencing the optimization of thermal fitness in warm-evolved mosquitoes, predicting that selection was strongest on the performance of juvenile development, followed by longevity and fecundity (Table 1, Supplementary Table 15). The identification of egg-to-adult viability as the dominant component of R₀ further redirects attention from adult-targeted interventions towards juvenile stages for mosquito population control under global warming^67^. We also found that the fitness trajectories of egg hatching time, egg size, larval viability, PDT, pupation rate and sex ratio did not change during acclimation and warm evolution, revealing that these traits are not thermally sensitive under our tropical regime. These results support the conclusion that mosquito traits respond differently to temperature, which cautions against using static and uniform parameterization of trait dependencies on T*_a_* in models of mosquito population dynamics under climate change^82–84^.

Finally, our results highlight a significant mismatch in thermal sensitivity between sexes, as reported across plants and animals^85–88^. Males showed a slower pace of phenotypic adjustment to tropical conditions than females. Sex differences in acclimation capacity are expected across ectotherms^89^ and we recently showed that *Ae. albopictus* males are more sensitive to heat stress than females^90^. We propose that the observed higher thermal resilience of females relative to males depends on the reproductive behavior and physiology of mosquitoes. *Aedes* spp. mosquitoes mate in small swarms, which are formed by several males trying to mate first with a female (scramble competition)^91,92^. Males can mate multiple times, while females are monandrous and require a blood meal from a vertebrate host to complete egg maturation^91–93^. Consequently, females tend to exhibit more exploratory behaviors and can be more aggressive than males, resulting in a faster pace of life and exposure to greater temperature fluctuations than males^94^. These findings also underscore the need to further investigate the thermal dependency of male physiology, because all current innovative control strategies targeting adult mosquitoes, such as the “sterile insect technique” and *Wolbachia*-based interventions, depend on the release of males^95–97^.

## Methods

### Experimental evolution with *Aedes albopictus*

We established ten replicate colonies, each starting with 1,000 *Aedes albopictus* eggs, as described in the results section ‘Thermal fitness of control mosquitoes remains stable during EE’. The microbiota of Foshan mosquitoes is known to be very stable under the same rearing conditions^98^, minimizing the risk of changes in the microbiota across our replicate lines. Larvae were reared in BugDorm plastic pans (19×19×6cm) and fed daily with 10 pellets of Tetra Goldfish Gold Colour fish food (Tetra Werke, Germany) until pupation. Adults were housed in cardboard cages (∼22 L of volume), with *ad libitum* access to 10% sucrose solution. In ectotherms, metabolic demands are expected to increase with temperature due to the acceleration in biochemical reactions^45^, conditioning effective per capita resource availability. To reduce fitness bias due to temperature-by-resource interactions, we provided food in excess during juvenile development.

We counted the collected pupae daily to maintain adult populations of around 800-900 individuals per cage. Seven to ten days post emergence, we offered defibrinated mutton blood (Biolife Italiana) to females using a Hemotek feeding apparatus. Females were offered five blood meals at each generation to collect eggs for the next generation. We mixed eggs from the second and third egg collection for the next generation to avoid selection for early or late fecundity. Mosquitoes were sampled for fitness, physiological and transcriptomic assessments at successive generations, for instance G_5_ and G_7_ or G_13_ and G_15_, to avoid creating artificial bottlenecks on lines, which could bias results. Consistent fitness values of control mosquitoes (Supplementary Figure 3) demonstrate the quality of our conditions and support biological conclusions on warm-evolved and relaxed-evolved mosquitoes.

### Mosquito fitness

We assessed *mosquito fitness* (*i.e.*, the contribution of an individual or genotype to the gene pool of subsequent generations^99^) through its major components: survival and reproduction. We traced changes in lifespan and developmental traits as a proxy of survival fitness; we measured the number of eggs during oviposition (fecundity), the number of hatching eggs (fertility) and that of larvae (progeny per female) as a proxy of reproductive fitness. We also measured female wing length, which is often used as a proxy for female fecundity^100,101^. Mosquito fitness was assessed for each replicate line every five generations during EE. We also integrated reproductive and survival traits into a fitness model to quantify the net female reproductive rate (R_0_) across generations as a proxy of ‘lifetime reproductive success’, see details in the corresponding section in Methods.

#### Reproductive fitness

To assess fecundity, five days post-emergence, females were offered a blood meal using an artificial membrane feeding system for 90 minutes starting at 3 PM. Between 12 and 20 fully engorged females per replicate were then carefully removed from the cage and placed individually into paper cups. Two days post-blood meal, egg-laying papers were provided, and oviposition was monitored for each mosquito. Eggs were hatched to determine fertility and number of resulting larvae monitored to quantify larvae/female. After oviposition, females were collected for wing length measurement as previously described^17^. Data distribution was tested for normality to decide whether to apply parametric ANOVA or non-parametric Mann–Whitney or Kruskal–Wallis tests to test for significant differences. Statistical differences were considered significant at P-value ≤ 0.05. Differences in fitness parameters between generations and across temperatures were analyzed using one-way ANOVA with Dunn’s multiple comparisons test. The Mann–Whitney test was used to compare fitness parameters between C.G_1_ and T.G_1_ during acclimation.

#### Developmental fitness

We measured egg hatchability using 150-200 eggs for each replicate colony. These eggs were placed in BugDorm plastic containers (17 x 6.5 x 12 cm) with 200 ml autoclaved water. Larval and pupal developmental time were calculated starting with a minimum of 100 first instar larvae for each replicate colony and following development of juveniles as previously described^17^. We collected pupae daily and transferred them to plastic cups filled with water from their original tray to allow for adult emergence. After emergence, we recorded the number of viable adults to assess “adult mortality” one day post-emergence. Adults were then sexed and Male Emergence Time (MET) and Female Emergence Time (FET) were calculated from the day of hatching to the day of adult emergence. Males and females were placed separately into labeled plastic cups according to their date of emergence. Adults that emerged on the same day were grouped together. A sugar-soaked cotton ball was placed on top of the mesh covering each cup, with 10% sucrose solution refreshed daily and the cotton replaced weekly. To assess adult longevity, adult cups were checked daily, and the number of dead individuals was recorded until all mosquitoes had died. Longevity data were used to estimate median survival time using the Mantel–Haenszel method^102^. Comparisons of mosquito longevity were performed using Kaplan–Meier survival analysis with the log-rank (Mantel–Cox) test^103^. To compare the temporal profile of daily adult emergence between sexes, we fitted cubic-spline models to the cumulative daily emergence of males and females for each thermal regime and generation. Best-fit parameters with 95% confidence intervals are reported in Supplementary Table 9, and fitted curves with survival probabilities are shown in Supplementary Figure 7. Differences in median lifespan across generations and temperatures were assessed with one-way ANOVA followed by Tukey’s multiple comparisons test. All statistical analyses were conducted using Prism 10 (GraphPad Software, San Diego, CA, USA).

#### Trait trajectories

We rendered each life-history and morphological trait as a continuous trajectory of percentage change from the control mean across generations and thermal regimes (Fig. 1B and C). For each trait, we expressed each condition as [100 × (mean_X_ − mean_control_) / mean_control]_, where mean_X_ is the trait mean at a given condition and mean_control_ is the mean value of controls. The sign of the metric was retained, so that positive and negative deviations from control remained distinguishable, and 0% marks the control value. For all fitness traits, the control reference was the mean of the three control generations (C.G_1_, C.G_5_, C.G_10_); wing length, which was measured only at the founder generation, was referenced to C.G_1_ alone. Lifespan was summarized as the median survival time of each replicate line. A grey band spanning ±10% marks the range within which trait values are treated as equivalent to control. We connected the trajectories of warm-evolved and the relaxed-lines of the same trait with a dotted segment linking the last tropical generation (T.G_20_) to the first relaxed generation (R.G_1_), so that each trajectory reads continuously based on the correspondence between warm-evolved and relaxed lines (Fig. 1A). Significance was taken from the post-hoc comparisons described above (Dunn’s or Tukey’s tests). Trajectories were computed and plotted in R (v4.5.0) with the tidyverse, ggplot2, patchwork and ggh4x packages.

#### Trajectory model

To place traits measured on different scales on a common axis, each trait mean was expressed as a standardized deviation from a pooled control reference, d = (mean_X - mean_control) / SD_pooled_control, where the reference pools the control lines at generations 1, 5 and 10 and SD_pooled_control is the ANOVA-style total standard deviation across the three control groups. Sample sizes were back-computed from the reported SD and SEM. Traits with data from less than three tropical generations were excluded, leaving 15 traits with complete generational coverage (generations 1, 5, 10, 15 and 20).

The magnitude of departure from control, |d|, was modelled on tropical generation in a linear mixed model with trait as a random intercept (statsmodels MixedLM), so that each trait contributes its own baseline, but a single generation term is estimated across traits. A quadratic generation term was added and compared with the linear model by likelihood-ratio test on maximum-likelihood fits. Per-trait slopes were also fitted individually and corrected across traits with the Benjamini–Hochberg procedure.

Traits were grouped by trajectory shape rather than magnitude: each trait’s profile of d across tropical generations 1–20 was mean-centred and traits were clustered by correlation distance with hierarchical average linkage. The number of clusters was chosen by mean silhouette width over k = 2–5. Cluster assignment robustness was assessed by 1,000 bootstrap resamples of traits, recording for each trait the proportion of resamples in which it co-clustered with its assigned partners. Divergence of the two resulting time courses was tested in a linear mixed model of signed d on generation × cluster with trait as a random intercept, and the interaction was assessed by likelihood-ratio test on maximum-likelihood fits.

### Phenotyping of physiological traits

We quantified mosquito energetic budget following a modified existing protocol^104^. Briefly, we measured total protein, carbohydrate, glycogen, total lipid, and triglyceride contents in females at emergence using a colorimetric protocol. We weighed individual females on a microbalance (Mettler AC100) before homogenization in 2 mL Eppendorf tubes containing 180 μL of lysis buffer (100 mM KH₂PO₄, 1 mM DTT, 1 mM EDTA, pH 7.4). Samples were centrifuged at 180 × *g* for 5 min at 4 °C and the supernatant was used for biochemical assays. Protein content was measured using the Bradford assay reagent (Sigma-Aldrich, B6916). A 2.5 μL aliquot of supernatant and 2.5 μL of lysis buffer were mixed with 250 μL of reagent in 96-well plates and read at 595 nm (CLARIOstar plate reader). BSA served as the standard. Total carbohydrates were extracted by adding 2.5 μL lysis buffer, 20 μL of 20% Na₂SO₄, and 1.5 mL of chloroform–methanol (1:2) to the homogenate, followed by centrifugation (180 × *g*, 15 min, 4 °C). Supernatants were reacted with the anthrone reagent (1.42 g L⁻¹ in 70% H₂SO₄), heated at 90 °C for 15 min, and read at 625 nm. Glycogen was extracted from the pellet via two washes with 80% methanol, then processed with anthrone as above. D-glucose was used as the standard.

Lipid and triglyceride contents were determined using the vanillin–phosphoric acid method, with neutral lipids (triglycerides) separated from polar lipids using silicic acid. For total lipid quantification, a 150 μL aliquot of the supernatant was evaporated at 90 °C, treated with 10 μL H₂SO₄, incubated (90 °C, 2 min), cooled (0°C, 5 min), and reacted with 190 μL of vanillin reagent (1.2 g L⁻¹ in 68% H₃PO₄), with absorbance read at 525 nm. For triglycerides, 500 μL of supernatant was evaporated at 90 °C, re-dissolved in 1 mL chloroform, treated with 200 mg silicic acid (Sigma) to bind polar lipids, centrifuged, and assayed as above. Triolein was used as the standard. Between 12-20 female mosquitoes were analyzed per replicate/temperature/generation. Energy reserve data were analyzed using one-way ANOVA followed by Tukey’s multiple comparisons test. All statistical analyses and figures were generated using Prism 10.

### Phenotyping of behavioral thermal traits

We assessed mosquito thermal traits in terms of KDT and T*_p_*, following standardized protocols^105,106^. To assess T*_p_* and KDT, 4-6-day-old sugar-fed females were collected in the morning, and experiments were conducted in the afternoon. T*_p_* was quantified using a custom-built aluminum thermal gradient plate (15.3–41.5 °C; gradient increments: 0.79 ± 0.29 °C). The gradient was generated by connecting one end of the plate to a cool-water bath and the other to a warm-water bath. Groups of 10 females from the same replicate/condition were released at the center of the plate and allowed a 5-minute acclimation period, followed by 30 minutes of free movement. The final resting position of each mosquito was recorded as its preferred temperature. A minimum number of 20 female mosquitoes were analyzed per replicate, temperature, generation. KDT was measured using a custom-built aluminum plate containing nine individual wells. Six-day-old sugar-fed females were placed in the wells at 25 °C, and temperature was gradually increased at a rate of 0.5 °C/min up to 50 °C. A Logitech C922 Pro camera positioned above the device recorded mosquito movement. The temperature at which movement ceased was scored as KDT. At least nine mosquitoes per replicate, temperature and generation were tested.

### Quantification of egg freezing point

Egg freezing point was determined using individual eggs. First, eggs were assessed using a microscope and selected for the assay if they were not hatched or collapsed. Each egg was affixed to a fine thermocouple using a minimal amount of thermal grease applied to the egg cuticle to ensure optimal thermal contact. The egg was then placed on an aluminum plate mounted on a Peltier element, which allowed for controlled and gradual cooling. Temperature data were recorded continuously using a thermocouple connected to a voltage input data acquisition module (Omega Engineering, CT, USA). As the temperature decreased, freezing was detected by a sudden spike in temperature, which was recorded as the egg freezing point. At least 12 eggs per replicate, temperature, and generation were analyzed. Thermal behavioral trait data were analyzed using one-way ANOVA. The Kruskal-Wallis test was applied to compare differences across generations and temperature treatments. All statistical analyses and figure generation were performed using Prism 10.

### Egg morphometric

Newly laid *Ae. albopictus* eggs were collected from each replicate of the C.G_5_, T.G_20_, and R.G_5_ condition after three days of desiccation following oviposition. Eggs were photographed under an AMSCOPE microscope (equipped with a MU1003 digital camera) using the AMSCOPE associated software. Egg morphometric measurements were performed on the acquired images after calibration, with pixel measurements converted to millimetres. Egg length was defined as the linear distance between the micropyle and the opposite end of the egg, whereas egg width was measured as the maximum distance perpendicular to the longitudinal axis. The egg index was calculated as the ratio of egg length to egg width. For each replicate line, 30 eggs were randomly selected for morphometric analysis. Differences in egg length, width, and egg index among the C.G_5_, T.G_20_, and R.G_5_ groups were analyzed using a Kruskal-Wallis test followed by Dunn’s multiple comparisons test in GraphPad Prism 10.

### Gene expression analyses

Many tissues and developmental stages may be thermally responsive. We chose to track gene expression of the fat body to match our energy reserve analyses. We dissected fat bodies from newly emerged females between 17:00 and 19:00 to minimize circadian variation^107^. For each experimental group, we prepared two biological replicates per replicate line, each consisting of a pool of 20 females. We sequenced a total of 36 RNA-seq libraries: four control lines at G₁, four tropical lines at G₁ and G₁₅, and three relaxed selection lines at G₁ and G₇. Warm-evolved lines were sampled at G_15_ for transcriptomic analyses because most fitness traits of warm-evolved lines (*i.e.* LDT, FET, MET, fecundity, fertility, progeny per female) assumed values not significantly different from those of controls starting at G_10_-_15_. RNA was extracted using the TRIzol Plus RNA Purification Kit (Invitrogen) and sent to BGI Genomics for strand-specific mRNA library preparation (DNBSEQ Eukaryotic Strand-specific mRNA library prep kit) and 2 × 150 bp paired-end sequencing on the DNBSeq platform. Each library yielded ≥ 24 million read pairs.

Raw reads were processed with the nf-core/rnaseq pipeline^108^ (v3.20.0; Nextflow v25.04.6) using default parameters: quality control with FastQC (v0.12.1), adapter trimming with Trim Galore (v0.6.10), alignment to the *Ae. albopictus* AalbF5 genome (GCF_035046485.1) with STAR (v2.7.11b), and gene-level quantification with Salmon (v1.10.3). Alignments were sorted with Samtools (v1.21) and evaluated with RSeQC (v5.0.2). All transcriptomic analyses used R (v4.4.3). Key packages: DESeq2 v1.34.0, WGCNA v1.72-5, topGO v2.54.0, GOSemSim v2.28.1, rrvgo v1.14.2, ComplexUpset, ggplot2, igraph, ggraph, graphlayouts, pheatmap, cowplot.

### Differential expression analysis

We performed differential expression analysis using DESeq2 (v1.34.0) via the nf-core/differentialabundance pipeline (v1.5.0). Ten pairwise comparisons were conducted: T.G₁ vs C.G₁ (acclimation); T.G₁₅ vs C.G₁ and T.G₁₅ vs T.G₁ (warm evolution); and seven relaxed selection contrasts (R.G₁ vs C.G₁, R.G₁ vs T.G₁, R.G₁ vs T.G₁₅, R.G₇ vs C.G₁, R.G₇ vs T.G₁, R.G₇ vs T.G₁₅, R.G₇ vs R.G₁). DEGs were defined as genes with |log₂ fold change| > 1 and Benjamini–Hochberg adjusted *P* < 0.01, after excluding genes with mean normalized counts < 5. A post-hoc outlier filter was used to flag genes (https://github.com/UmbertoPalatini/DeSeq2_DEGs_OutlierDetector) in which a single sample contributed >40% of the log₂ fold-change magnitude, using ratio-based (>20 × median), IQR (3 × IQR), and extreme-value (>1,000 × minimum) thresholds; flagged genes were excluded unless fold changes remained consistent after removing the outlier sample. Volcano plots and per-comparison DEG counts derive from this full output; however, for grouped trajectory analyses, we strictly excluded genes with mean normalized counts < 5 to preserve statistical power.

### Transcriptomic similarity and expression trajectories

We assessed overall transcriptomic similarity among samples by Principal Component Analysis (PCA) and hierarchical clustering. PCA was computed with prcomp on the variance-stabilizing-transformed (VST) count matrix (DESeq2), with genes scaled to unit variance, and samples were projected onto the first two principal components (Fig. 2B). Pairwise sample distances were computed as Pearson correlation distance (1 − *r*) on log₂-transformed normalized counts, excluding genes with mean normalized count < 5, and displayed as a heatmap ordered by complete-linkage hierarchical clustering (Supplementary Figure 12). To quantify divergence from the control baseline, we computed the Euclidean distance between each replicate line and the C.G₁ centroid. Normalized counts were log_2_-transformed (log_2_[count + 1]) and averaged within each line (mean of the two pools); the C.G_1_ centroid was the mean across the four control lines over all genes with mean normalized count ≥ 5 (Fig. 3A).

Shared and unique DEG sets across contrasts were compared using directional UpSet plots (ComplexUpset R package), classifying genes as concordant or discordant by direction of change. For warm evolution, the three comparisons of T.G_1_ vs C.G_1_; T.G_15_ vs C.G₁; and T.G_15_ vs T.G_1_ were intersected; for relaxed selection, the lists of genes from the comparisons T.G₁ vs C.G₁ and R.G₁ vs T.G₁₅ were intersected using the same thresholds and directional classification.

### Reversal score

To quantify transcriptional reversal upon return to control conditions, we computed a per-gene reversal score as:

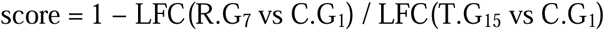

where LFC values are DESeq2 log₂ fold changes, both expressed relative to the same reference group (C.G_1_). Because both LFC values share the same reference, the ratio is sign-consistent for each gene regardless of the direction of regulation: a gene upregulated in T.G₁₅ relative to C.G_1_ retains a positive LFC in R.G_7_ vs C.G_1_ if expression remains elevated, and a negative LFC if it drops below control. Negative scores, which would arise if R.G₇ expression changed in the direction opposite to T.G₁₅, were not observed. The score was computed independently for each gene from its own LFC estimates.

Genes were retained if they were significant DEGs between T.G_15_ vs C.G_1_, had |LFC(T.G_15_ vs C.G_1_)| > 0.3 to stabilize the ratio against noise at small fold changes, and had mean normalized counts ≥ 10 (a stricter threshold than the ≥ 5 cutoff used for ever-DEG classification, given the ratio’s sensitivity to estimation noise at low expression).

### Weighted gene co-expression network analysis

We used the R WGCNA package (v1.72-5)^109^ separately for acclimation (C.G₁ vs T.G₁) and warm evolution (C.G₁, T.G₁, T.G₁₅) samples. After excluding outlier-flagged genes and those with mean expression < 5, we retained genes in the top 60th percentile of expression variance. Signed hybrid networks were constructed using a soft-thresholding power selected via the pickSoftThreshold() function (WGCNA v1.72-5), which evaluates scale-free topology fit (R²) across a range of candidate powers. We selected the lowest power at which the signed R² exceeded 0.85, corresponding to a soft-thresholding power of 10 for the acclimation network and 6 for the warm evolution network. Modules were identified by hierarchical clustering of the TOM dissimilarity matrix (average linkage, minimum module size = 30, deep split = 2); modules with eigengene correlation ≥ 0.70 (merge cut height = 0.30) were merged. Module eigengene differences were tested by two-sample *t*-tests (acclimation) or one-way ANOVA with pairwise *t*-tests (adaptation), with Benjamini–Hochberg correction (FDR < 0.01). Hub genes were the top 10 per module by intramodular connectivity (kWithin).

### Gene ontology enrichment and GO Slim categorization

GO enrichment was performed with topGO (v2.54.0; weight01 algorithm, Fisher’s exact test) for each WGCNA module, UpSet gene set, and reversal category, testing Biological Process (BP) and Molecular Function (MF) ontologies against a background of all detected genes (mean normalized count > 1). GO annotations were taken from the AalbF5 RefSeq Gene Association File (GCF_035046485.1). Terms with fewer than 5 or more than 500 annotated genes were excluded, and terms with P < 0.01 were retained as significant.

Redundancy among enriched GO terms was reduced in two steps. First, semantic similarity between pairs of GO terms was computed with the Wang method (GOSemSim v2.28.1); term information content was derived from *Drosophila melanogaster* (org.Dm.eg.db), which served as the reference OrgDb for GO graph weighting, as a well-annotated *Aedes*-specific OrgDb is currently unavailable. Because Wang semantic similarity is calculated from the structure of the GO hierarchy and the information content of shared ancestor terms, which are species-independent at the GO term level, the use of *D. melanogaster* information content does not affect the identity of the GO terms being compared; these terms were assigned to *Ae. albopictus* genes from the AalbF5 RefSeq Gene Association File as described above. Second, GO terms were clustered at a semantic similarity threshold ≥ 0.7 (rrvgo v1.14.2), and the most significant term per cluster was retained. For WGCNA modules, up to 30 non-redundant terms were displayed as networks (edges at similarity ≥ 0.15; node size = gene count; node color = −log₁₀ P; stress-majorization layout via graphlayouts). For reversal categories, up to 10 terms per category were shown.

For cross-comparison visualization, each significant GO term was assigned to a functional category by matching the term or its GO ancestors to a curated set of representative GO IDs, assigning each term to the first matching category and terms with no match to “Other.” The categories spanned metabolism, RNA processing, transcription, translation, protein processing, cell cycle, DNA repair, stress response, immune response, signaling, transport, development, cell death, cytoskeleton, cell adhesion, reproduction, and chromatin.

### Immune gene identification and analysis

A curated *Ae. albopictus* immune gene catalogue published with the AalbF3 assembly^110^ (661 entries) was remapped to the AalbF5 assembly (GCF_035046485.1). Each legacy identifier was resolved through NCBI Gene history and retained only if it corresponded to a current AalbF5 gene model present in the expression matrix; retired identifiers were merged into a single model, identifier that no longer returned a current gene were discarded. This yielded 360 immune genes assigned to 22 functional families (348 with a family assignment; family size 5-61 genes).

We first compared the distribution of DESeq2 Wald statistics for immune genes with that of all other tested genes using two-sided Wilcoxon rank-sum tests, on signed statistics (directional shift) and on absolute statistics (magnitude of response); 331-342 immune genes were testable per contrast after DESeq2 independent filtering. Second, normalized expression was examined directly: counts were log2-transformed, averaged within replicate line (the two sequencing pools per line were averaged first), and each gene was z-scored across the five conditions, so that zero denotes a gene’s own mean across the experiment. The mean immune-gene z-score per condition was compared with an expression-matched null of 4,000 random gene sets of the same size, matched to the immune set by decile of mean expression, which controls for the dependence of expression variability on expression level. Family-level tests applied the same Wilcoxon framework to each family against all other tested genes. P-values within each family of tests were corrected with the Benjamini–Hochberg procedure. Analyses used the same normalized count matrix and DESeq2 result tables as the transcriptome-wide analyses.

### Inter-replicate variance analyses

We quantified expression variance among independently evolving replicate lines, so that variance reflects divergence among lines under the same selection regime. Per-gene inter-replicate variance was computed across replicate-line means for each condition (C.G_1_, T.G_1_, T.G_15_, R.G_1_, R.G_7_; *n* = 4, 4, 4, 3, 3 lines). DESeq2-normalized counts were log₂-transformed (log₂[count + 1]) and averaged within each line (mean of the two pools). Variance was expressed as log₂(var + ε), with ε = 0.001; delta variance relative to C.G₁ was the difference in log₂ variance between each condition and C.G₁ (Fig. 3D).

Genes were grouped into five categories: maintained (*n* = 250), reversed (*n* = 961), over-reversed (*n* = 42), other ever-DEGs, and never-DEGs. Differences in delta variance between conditions within a category were tested by paired Wilcoxon signed-rank tests (paired by gene) with Benjamini–Hochberg correction (Fig. 3D).

To examine the relationship between expression level and inter-replicate variance, we plotted mean expression (log₂ normalized counts) against log₂(variance + ε) at T.G₁, T.G₁₅, R.G₁, and R.G₇ for three gene sets: maintained genes (M, *n* = 250), all other DEGs (D), and the non-DEG background (B) (Fig. 3E). Linear fits and Spearman’s rank correlation coefficient (ρ) were computed separately for each gene set within each condition. The non-DEG background provides the reference against which the maintained and other-DEG sets are compared because log-transformed count data carry an intrinsic negative mean–variance relationship. Therefore, we interpret the mean–variance relationship of maintained genes relative to this background rather than in absolute terms. Because maintained genes are defined from their R.G_7_ expression, we interpret the R.G_7_ mean–variance relationship accordingly. This analysis follows the framework of Lai & Schlötterer^37,38^, in which variance reduction without mean change after long-term experimental evolution indicates stabilizing selection and concurrent change in both indicates directional selection.

### Ecological modelling of “lifetime reproductive success”

We calculated synthetic demographic indicators across thermal treatments and generations to compare fitness outcomes. Specifically, we computed R₀ using a constrained life-table approach that reflects the feeding and oviposition schedule maintained during EE.

Following demographic theory^50,111^, we defined R₀, which is the average number of female offspring that a single adult female is expected to produce over her lifetime, as:

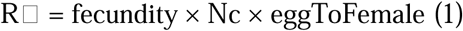

where ‘fecundity’ is the mean number of eggs laid per female per oviposition following the first blood meal. For each replicate group, we calculated fecundity as the total number of eggs collected divided by the number of blood-fed females. The term ‘Nc’ is the number of scheduled ovipositions a female can perform during her lifetime. The gonotrophic cycle is deduced based on our rearing schedule: females received their first blood meal 5 days after emergence and were blood-fed every 7 days. Assuming a fixed oviposition lag, L, of 2 days post-feeding, the age at first oviposition is:

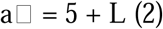

and hence the number of completed ovipositions is:

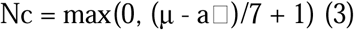

where ‘μ’ is the mean female longevity in days.

Finally, the ‘eggToFemale’ term is the probability that an egg results in an adult female, computed as:

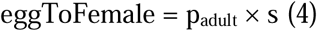

where ‘p_adult’_ is the egg-to-adult emergence rate and **s** is the observed female sex ratio among emerged adults. R₀ is valid under the assumptions that (i) all scheduled blood meals are successful, (ii) oviposition follows each feeding after lag L, (iii) fecundity remains constant across ovipositions, despite being only measured after the first blood meal, and (iv) female survival follows the mean longevity μ, with deterministic survival to each scheduled oviposition. Using fecundity measured after the first gonotrophic cycle as a proxy for data from subsequent ovipositions may lead to mild overestimation of lifetime fecundity. However, this approach provides a standardized, comparable fitness proxy across treatments, which is not biased by potential uneven effects of aging or blood meal on females from different treatments. Sensitivity to the oviposition lag was evaluated by varying L ∈ {1, 2, 3}. All the metrics were aggregated at the replicate × generation × treatment level, and R₀ was computed accordingly. All calculations were implemented in R (v4.5.0) functions from the *base* and *tidyverse* packages.

### Statistical modelling

To investigate temporal changes in fitness under thermal selection, we modeled R₀ across generations in the tropical treatment for oviposition lag equal to 2. We first fitted a univariate model where R₀ was regressed against generation number, including both linear and quadratic terms to capture potential non-linear trends in evolutionary trajectories. This model was specified as:

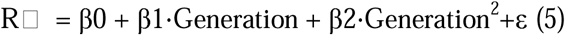

To identify the key life-history traits contributing to variation in R₀, we implemented a multivariate linear model incorporating standardized predictors. Specifically, we included fecundity, female longevity, and the probability of an egg developing into an adult female as explanatory variables. To account for temporal trends across the experiment, we also included generation as a covariate and captured potential non-linear dynamics by including a quadratic term (Generation²). Prior to model fitting, all continuous predictors were standardized (mean = 0, standard deviation = 1) to enable direct comparison of effect sizes across variables. The final model was specified as:

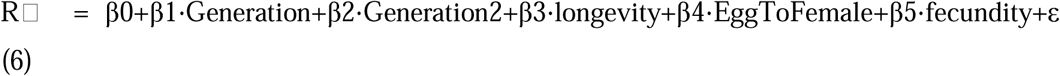

Model assumptions were verified through residual diagnostics, including visual inspection of residual plots and normality checks. To assess potential multicollinearity among predictors, we computed variance inflation factors (VIFs). As expected, VIFs for Generation and Generation² were elevated due to their mathematical dependence, but all other predictors exhibited acceptable VIF values (< 2.2). Given that the Generation terms were included to capture overall curvature rather than for individual interpretation, we retained them in the final model.

## Acknowledgements

We thank Claudio Lazzari and José Crespo for the KDT device design, Mélanie Body for designing the colorimetry protocol, Sandra Gabbert and Donald Mullins for support with egg freezing point measurements, Stefano Quaranta for initial handling of transcriptomic data and Elizabeth McGraw for helpful discussions.

## Author Contributions

A.K. maintained all mosquito lines, performed fitness and behavioral assessments, quantification of energy reserves, nucleic acid extractions and contributed to writing. U.P. performed all the bioinformatics data processing and downstream analyses, including differential expression, co-expression network analysis, GO enrichment, and variance trajectory analyses and contributed to writing. U.P. and A.K. prepared the final figures. D.D.R. conducted ecological analyses, modelling and contributed to writing. I.L-C. performed overall data analyses and data visualization, interpretation of results and drafted the final version of the manuscript. R.B., H.P. and S.D.C. contributed to mosquito maintenance and fitness assessments. R.R. contributed to ecological analyses, modelling and overall data analyses. H.A. contributed to fitness and behavioral experiments with A.K. M.B. conceived the study with C. L., secured funding, supervised the study, contributed to data analyses and drafted the final version of the manuscript.

## Data availability

The RNA-seq raw reads generated in this study have been deposited in the NCBI Sequence Read Archive under BioProject accession PRJNA1495493. All differential expression results are provided in Supplementary Tables 5, 10, 11, 16, 17, 20, and 21. All life-history, physiological, and behavioral measurements and statistics underlying Fig. 1 and Supplementary Figures 1 to 3 and 6 are provided in Supplementary Tables 1 to 4, 7, and 8.

## Code availability

The post-hoc DESeq2 outlier filter custom R script is available at https://github.com/UmbertoPalatini/DeSeq2_DEGs_OutlierDetector.

## Supplementary Figures

**Supplementary Figure 1.**
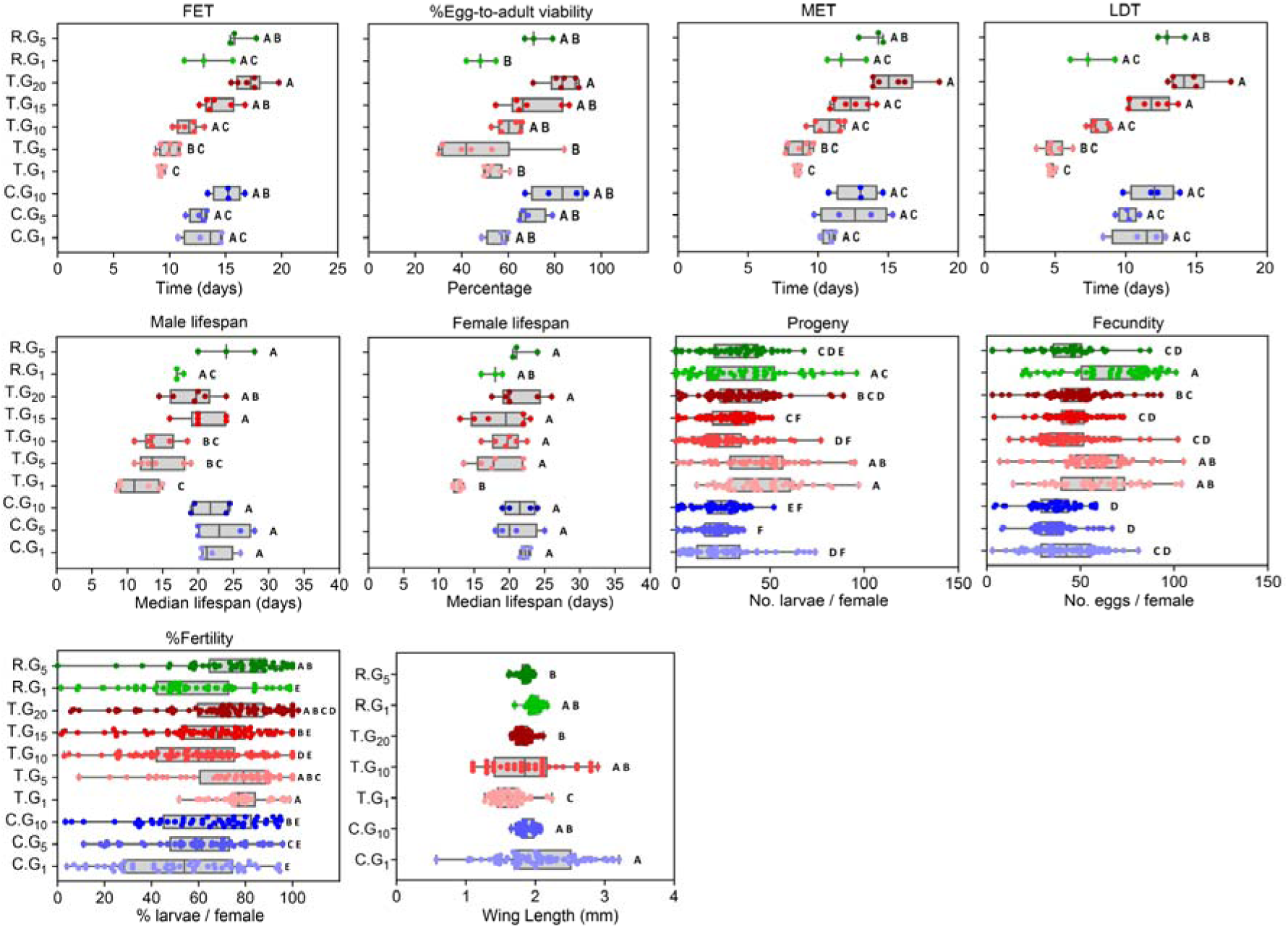
Life-history traits across thermal regimes and generations for thermally sensitive traits: female emergence time (FET), male emergence time (MET), larval developmental time (LDT), percentage egg-to-adult viability, male and female median lifespan, progeny per female, fecundity (eggs per female), percentage fertility (larvae per female), and wing length.

**Supplementary Figure 2.**
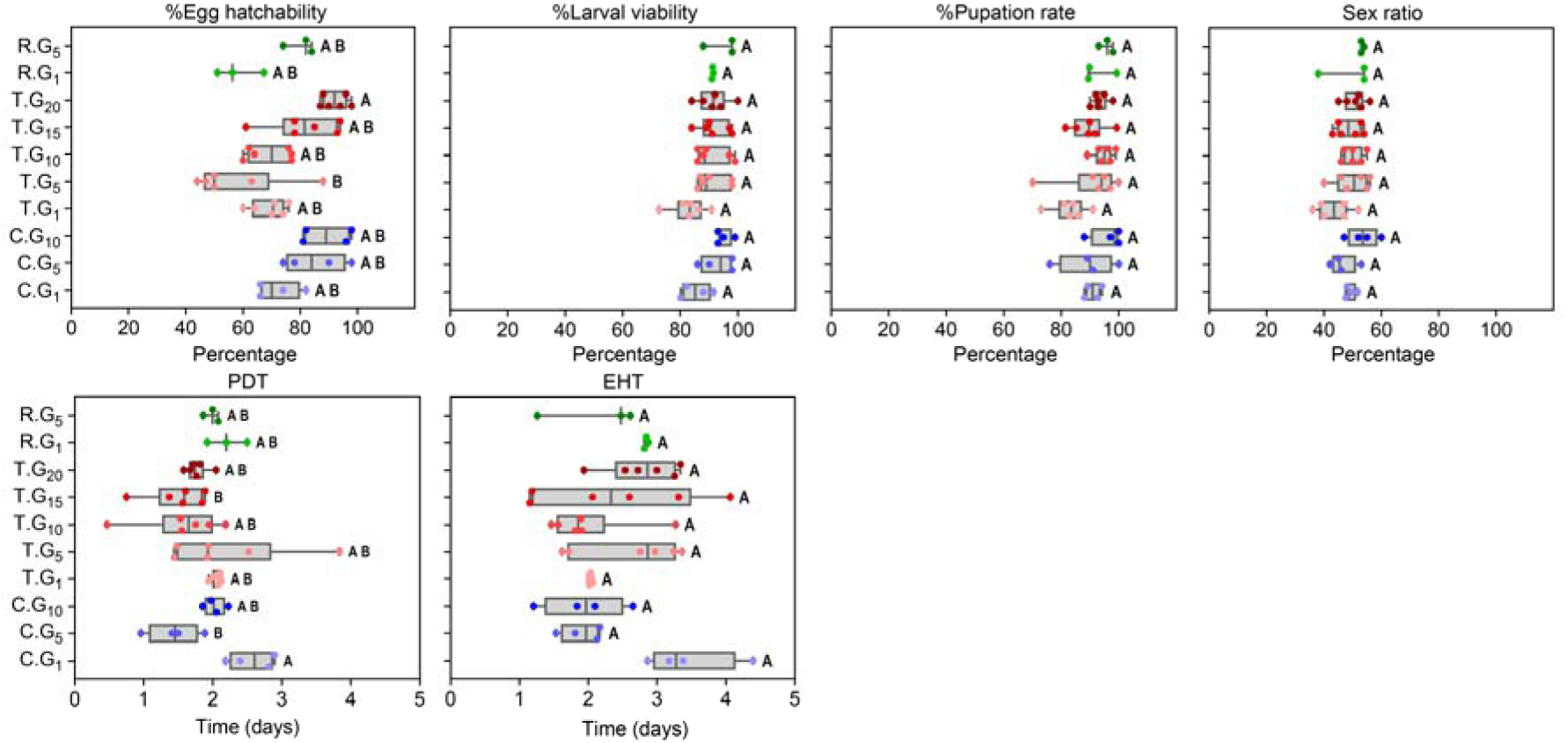
Life-history traits across thermal regimes and generations for thermally stable traits: percentage egg hatchability, percentage larval viability, percentage pupation rate, sex ratio (percentage females), egg hatching time (EHT), and pupal developmental time (PDT). Data are shown for control (C, blue), tropical (T, red), and relaxed-selection (R, green) mosquitoes at the indicated generations, with shading darkening with generation. Boxes span the interquartile range, the center line is the median, and whiskers extend to the most extreme values; individual measurements are overlaid. Letters denote compact letter display groupings; shared letters indicate no significant difference (α = 0.05). Sample sizes and statistical procedures are given in the supplementary materials and Supplementary Table 2.

**Supplementary Figure 3.**
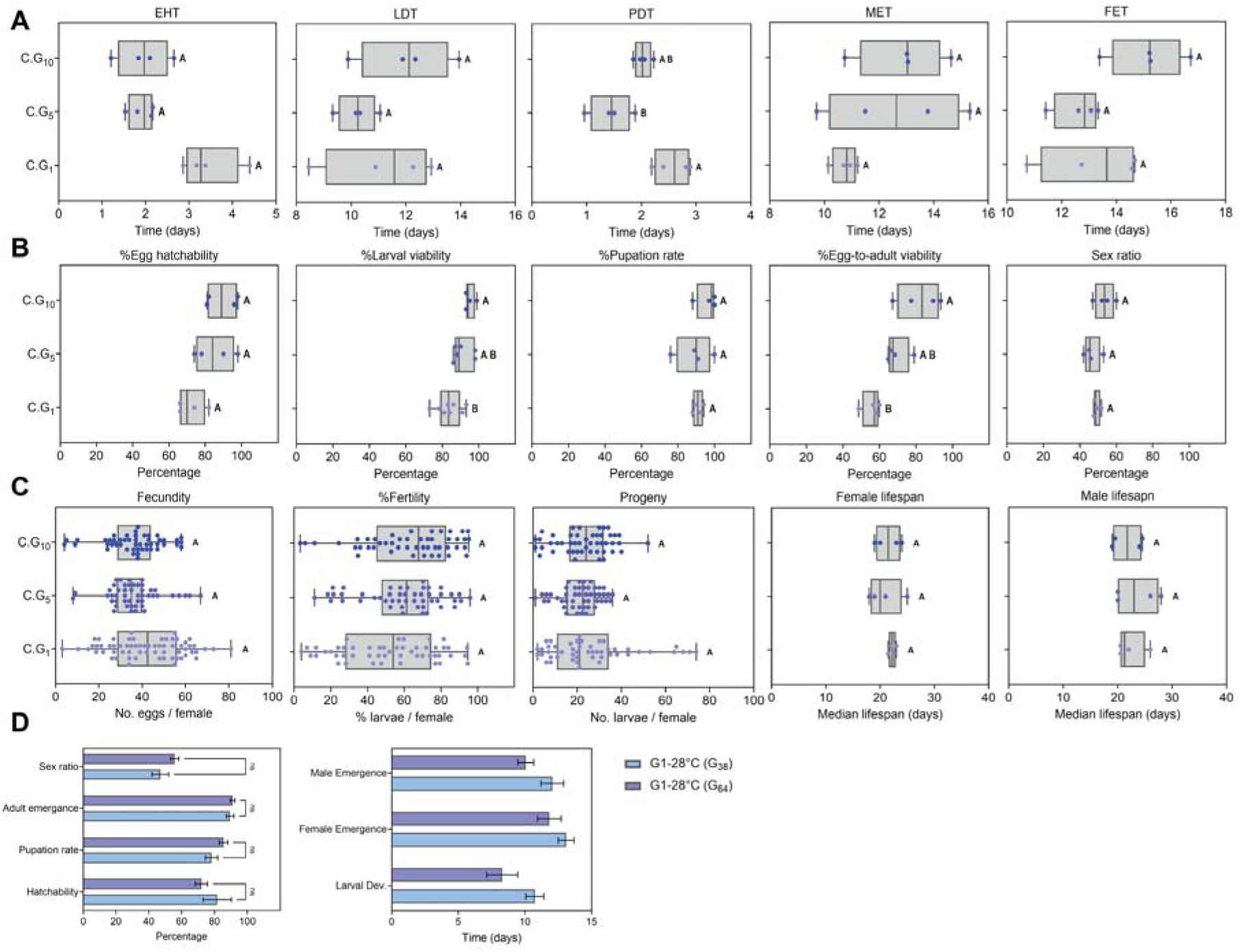
Comparison of fitness traits of control mosquitoes across generations (G) during and before EE. (**A**) Time (days, X axis) for EHT, LDT, PDT, FET, and MET. (**B**) Percentage (X axis) of egg hatchability, larval viability, pupation rate, egg-to-adult viability, and sex ratio. (**C**) Percentage (X axis) of fecundity, fertility, and progeny, and median lifespan (days, X axis) of females and males. All data are from mosquitoes maintained under control conditions (C); blue shading darkens with generation, from G₁ to G₅ and G₁₀. Letters denote compact letter display groupings; shared letters indicate no significant difference (α = 0.05). Sample sizes and statistical procedures for each trait are given in the Methods. (**D**) Fitness of Foshan mosquitoes at 38 generations (G₃₈) after establishment in the insectary at the University of Pavia, and at G₆₄, when EE started.

**Supplementary Figure 4.**
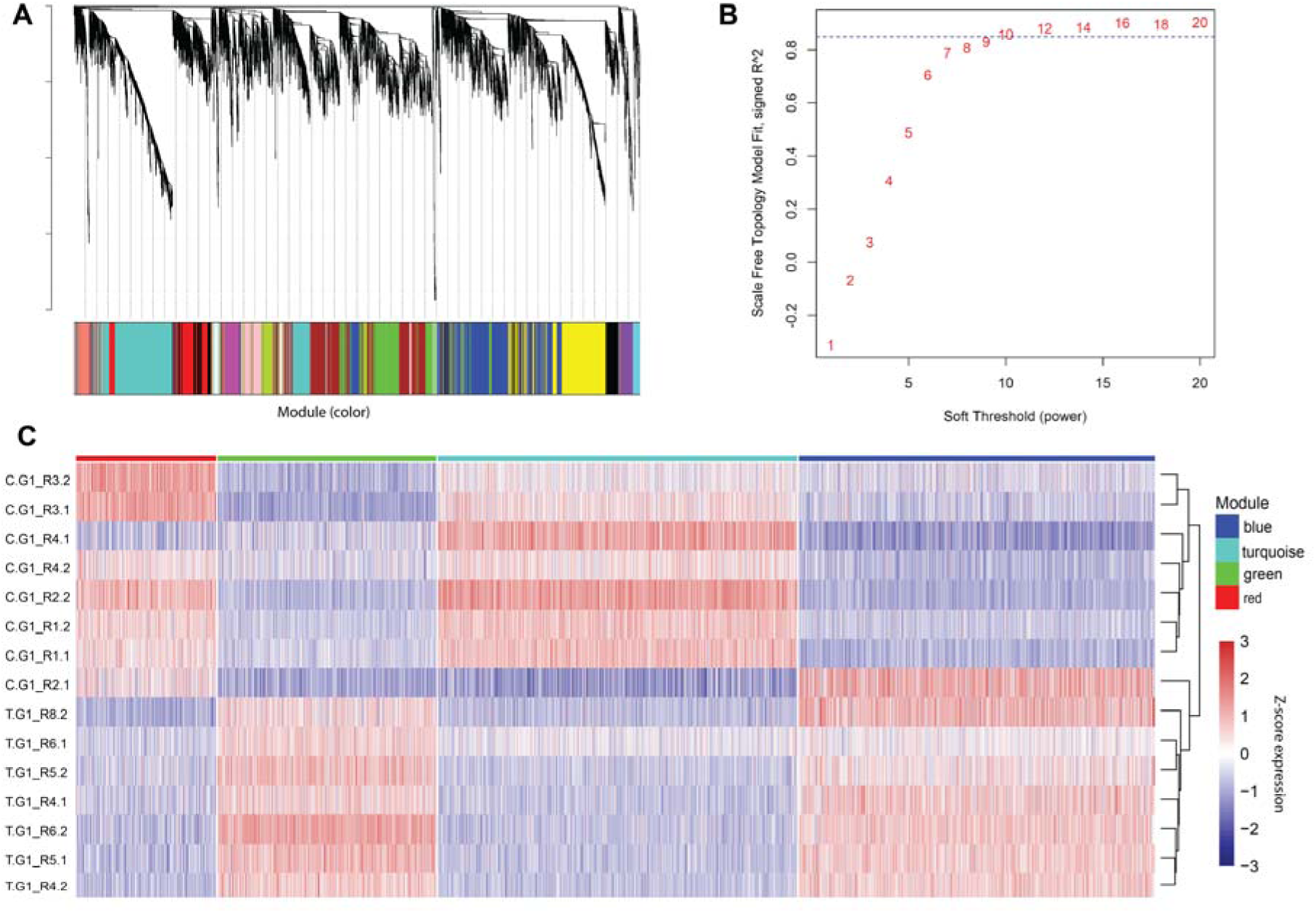
Co-expression network construction during thermal acclimation. (**A**) Gene dendrogram from hierarchical clustering of fat body transcriptomes (C.G₁ and T.G₁). The color band indicates module assignment by the WGCNA dynamic tree-cutting algorithm. (**B**) Scale-free topology model fit (signed *R*²) as a function of soft-thresholding power. The dashed line marks the *R*² = 0.85 threshold; a power of 10 was selected for network construction. (**C**) Heatmap of expression for genes assigned to the blue, red, green, and turquoise modules in C.G₁ and T.G₁ pools. Columns are individual sequencing pools (labeled by replicate line) clustered by expression profile; rows are genes. Colors indicate row-normalized *Z*-scores of normalized expression counts. Sample sizes and statistical procedures are given in the supplementary materials and Supplementary Table 6.

**Supplementary Figure 5.**
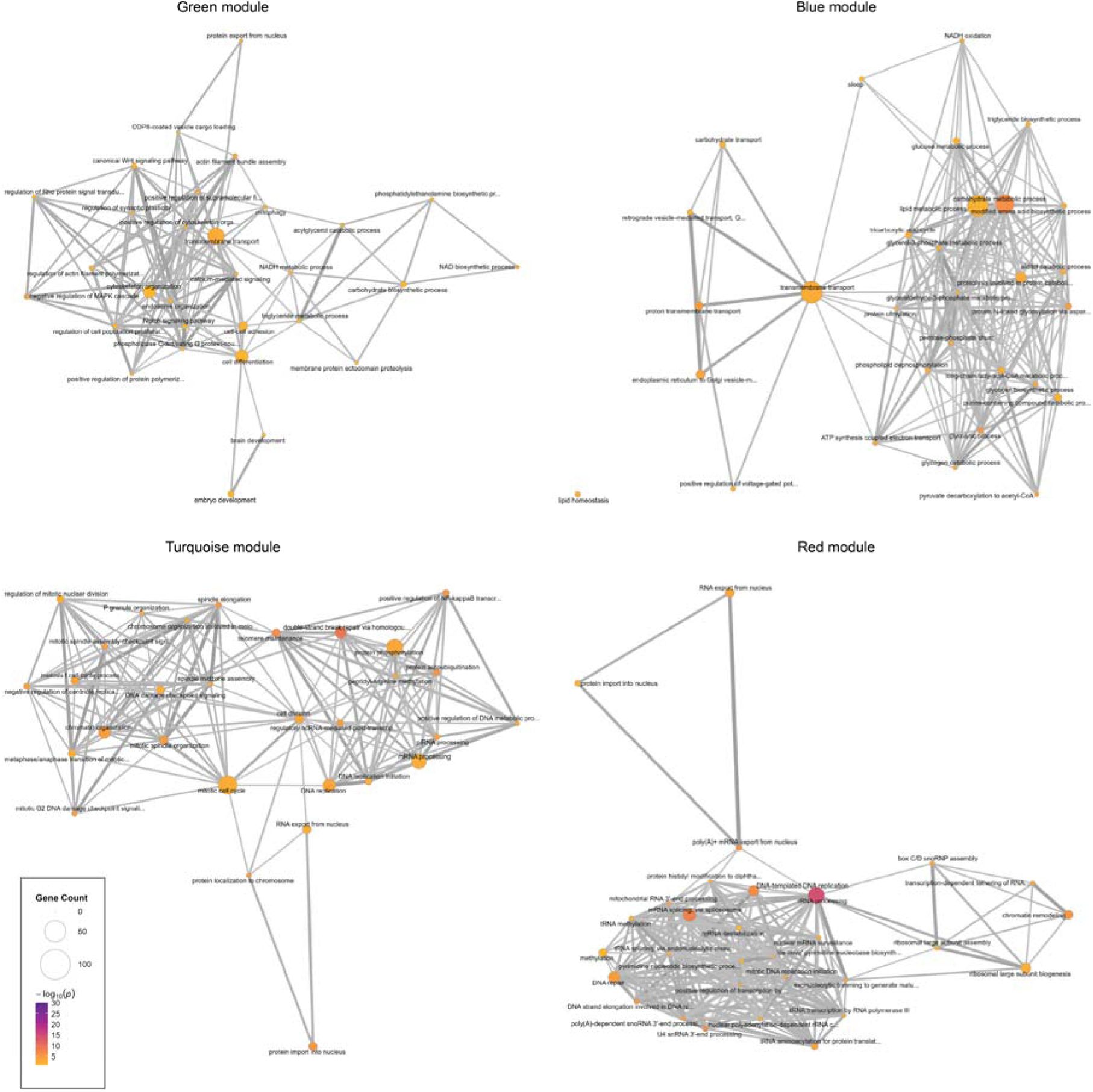
GO enrichment networks for the acclimation co-expression modules. Biological Process GO terms for the four primary WGCNA modules (green, blue, turquoise, red) identified in the C.G₁ versus T.G₁ comparison. Nodes are GO terms; node area is proportional to the number of module genes annotated to each term, and node color indicates enrichment significance (−log₁₀ *P*). Edges link terms with Wang semantic similarity ≥ 0.15 (GOSemSim), with edge width proportional to the similarity. Node area, node color, and edge width use a common scale across all panels. Redundant terms were collapsed at a semantic-similarity cutoff of 0.7 and the 30 most significant terms per module retained; layouts were computed by stress majorization. Full enrichment results are given in Supplementary Table 6.

**Supplementary Figure 6.**
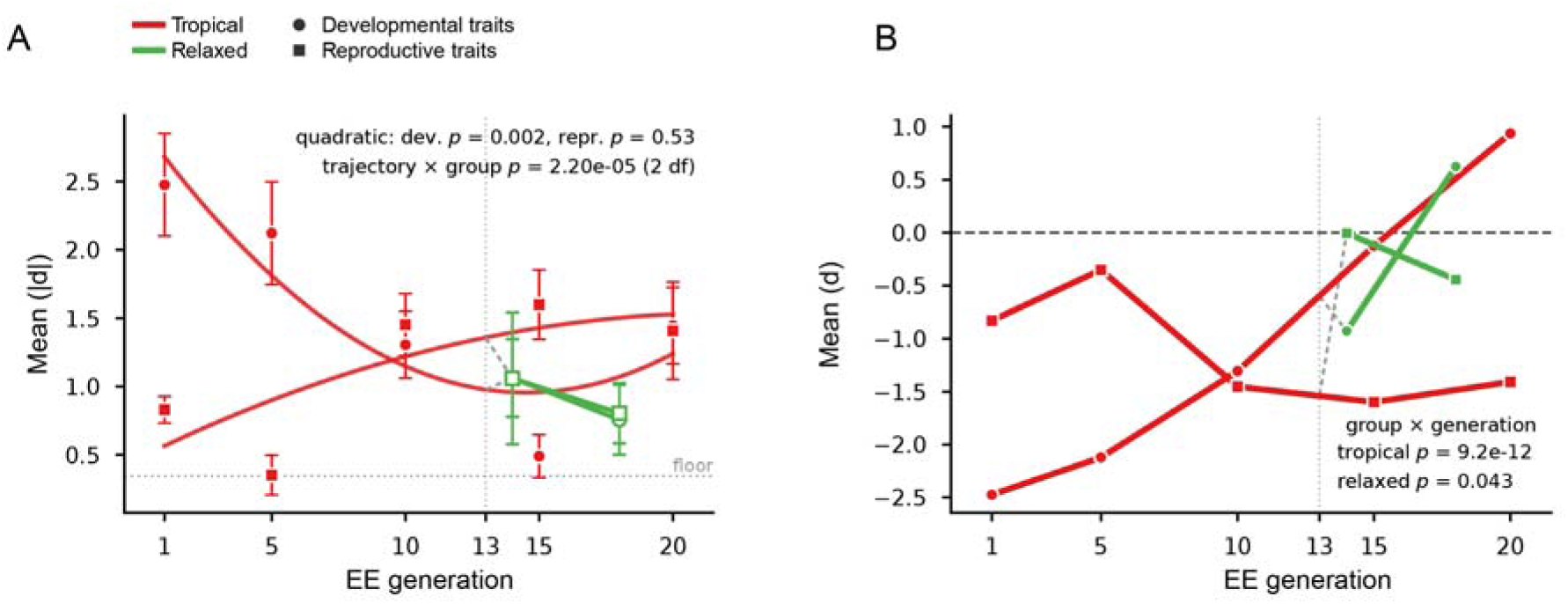
Fitness departure from control across tropical generations. (**A**) Mean absolute standardized deviation from the pooled control reference (|d|; control lines at generations 1, 5 and 10) across the 15 fitness and life-history traits with complete generational coverage. Points are the mean across traits ± s.e.m.; the curve is the fitted quadratic term of a linear mixed model with trait as a random intercept, and the dotted line marks its minimum at generation 14. P-values are for the quadratic coefficient and for the likelihood-ratio test of the quadratic against the linear model. (**B**) Standardized deviation from the pooled control reference (d; control lines at generations 1, 5 and 10) for each of 15 traits across tropical generations, coloured by trajectory cluster; thin lines are individual traits and thick lines the cluster means. Zero denotes the control value; the P-value is for the generation × cluster interaction in a linear mixed model with trait as a random intercept.

**Supplementary Figure 7.**
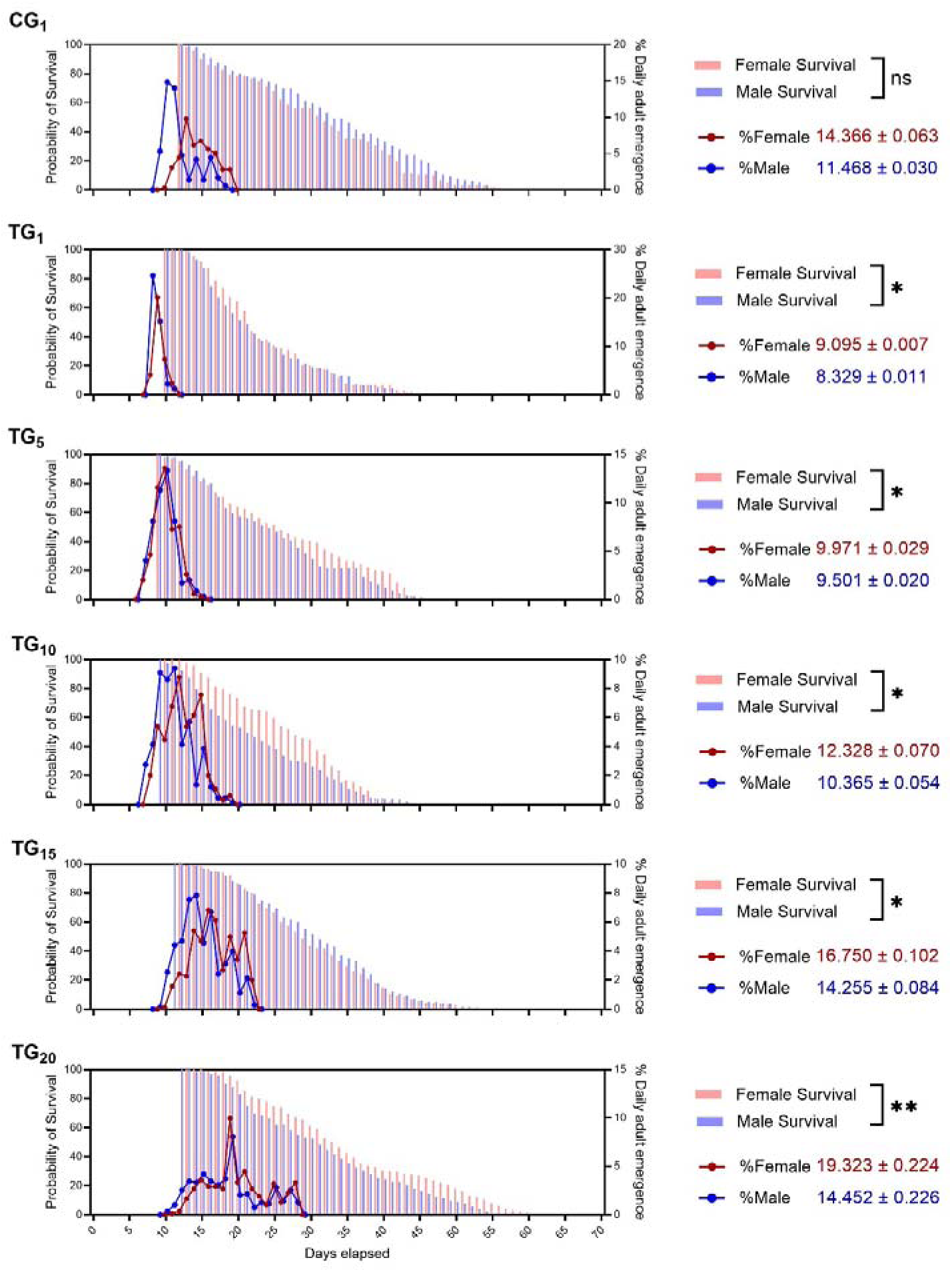
Shift between the percentage of daily emergence in females and males and respective longevity in control mosquitoes (C.G_1_) and warm-evolved (T) mosquitoes across generations (G). Each panel shows, on the Y axis, the probability of survival (left) and the percentage of daily emergence (right) of females (red) and males (blue) with respect to time, in days (X axis). Next to each plot, we report results of the cubic spline analysis assessing the statistical significance of the comparison between female and male survival time; significance levels are: ** p<0.01, *** p<0.001, **** p<0.0001. We also show the mean (and standard error) for the daily percentage of emergence time of females (red) and males (blue).

**Supplementary Figure 8.**
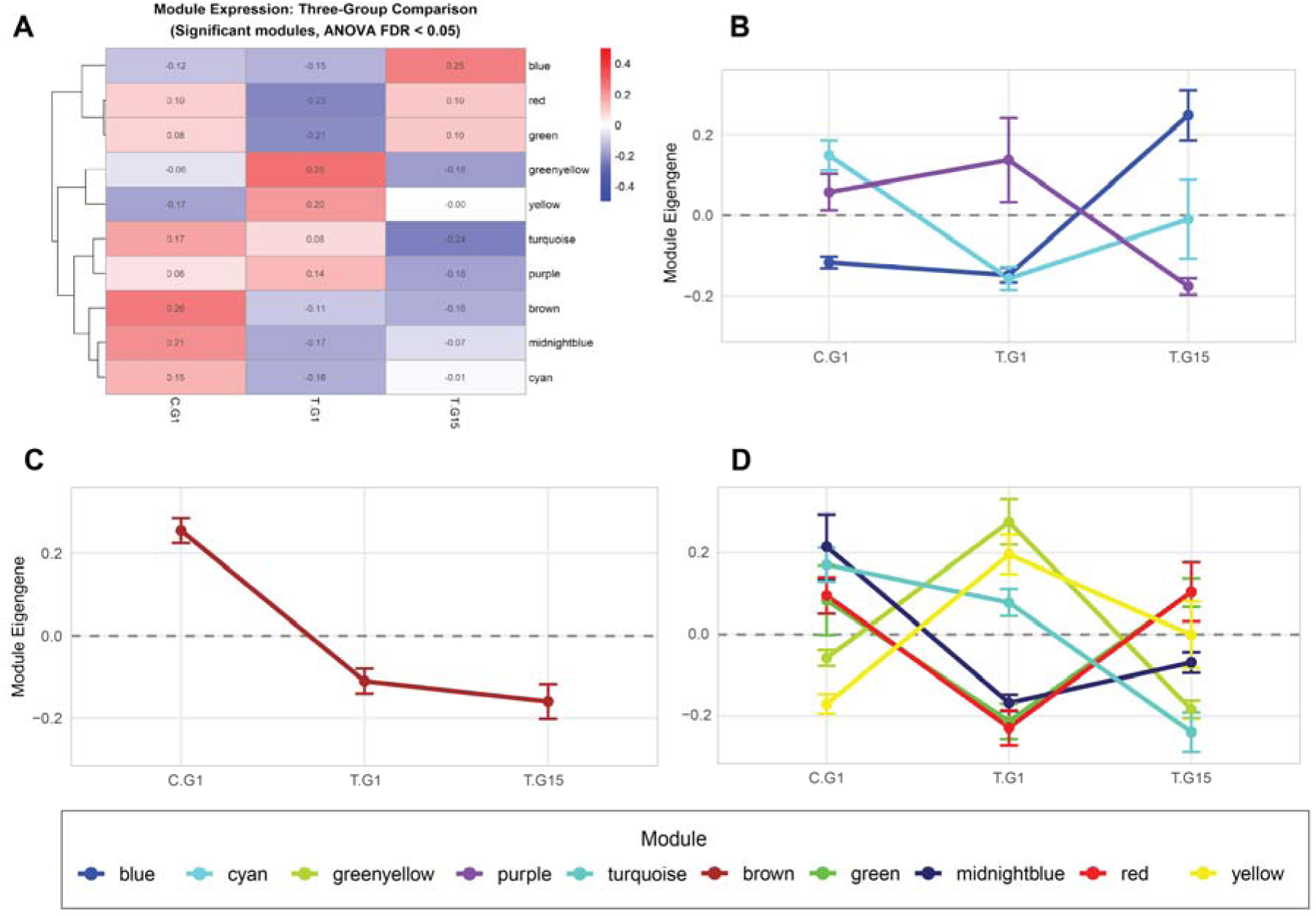
Module eigengene trajectories across acclimation and warm evolution. (**A**) Module eigengene values for the significant WGCNA modules (ANOVA FDR < 0.05) in control (C.G₁), warm-acclimated (T.G₁), and warm-evolved (T.G₁₅) mosquitoes. Cell color and value indicate the mean eigengene per group; rows are hierarchically clustered. (B to D) Module eigengene trajectories across the three conditions, grouped by expression pattern: modules peaking at T.G_15_ (**B**), modules highest at C.G₁ and declining thereafter (**C**), and modules peaking at T.G_1_ (**D**). Points are mean eigengene ± SEM; colors denote modules as in the key. The color band indicates module assignment; ten co-expression modules with significantly different expression patterns were identified.

**Supplementary Figure 9.**
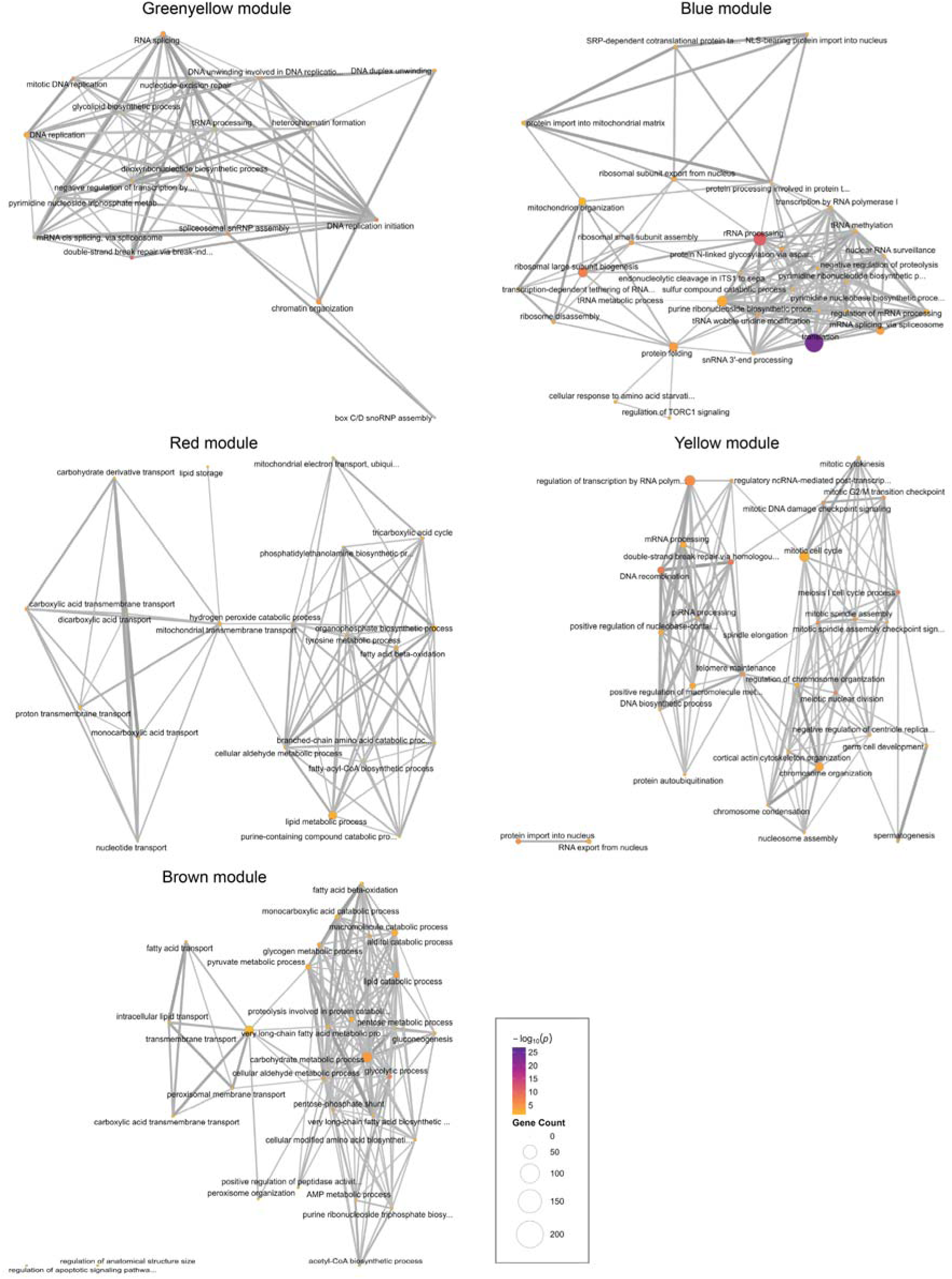
GO enrichment networks for the warm-evolution co-expression modules. Enriched Biological Process GO terms for the significant WGCNA modules identified in the three-group comparison (C.G₁, T.G₁, T.G₁₅), shown here for the greenyellow, blue, red, yellow, and brown modules. Nodes are GO terms; node area is proportional to the number of module genes annotated to each term, and node color indicates enrichment significance (−log₁₀ P). Edges link terms with Wang semantic similarity ≥ 0.15 (GOSemSim), with edge width proportional to the similarity. Node area, node color, and edge width use a common scale across all panels. Redundant terms were collapsed at a semantic-similarity cutoff of 0.7 and the 30 most significant terms per module retained; layouts were computed by stress majorization. Full enrichment results are given in Supplementary Table 14.

**Supplementary Figure 10.**
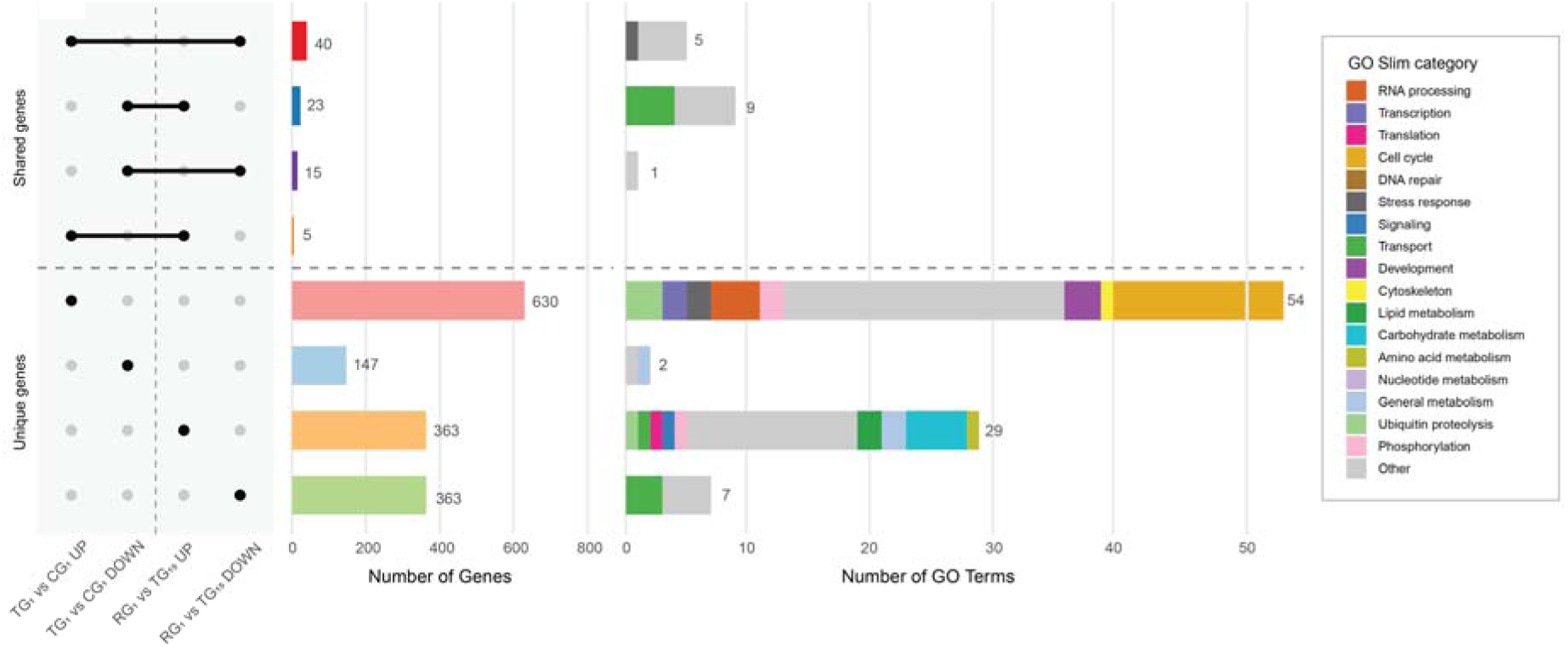
Comparison between acclimation signature of T.G_1_ and R.G_1_ mosquitoes. Directional UpSet plot of DEGs in the fat body, intersecting the acclimation contrast (T.G₁ vs C.G₁) with the return to control conditions (R.G₁ vs T.G₁₅). Each contrast is split by direction of regulation, giving the four sets on the *x* axis of the matrix (left). Filled dots mark the sets contributing to each intersection; the horizontal dashed line separates intersections shared between the two contrasts (top) from genes unique to one contrast (bottom). Left bars, number of genes per intersection; the 83 shared genes comprise 40 upregulated at T.G₁ and downregulated at R.G₁, 23 downregulated at T.G₁ and upregulated at R.G₁, 15 downregulated in both, and 5 upregulated in both, so 63 change in opposite directions and 20 in the same direction across the two transitions. Right bars, number of enriched Gene Ontology terms for the genes of each intersection, stacked by GO Slim functional category (key). DEGs were called with DESeq2 (|log₂ fold change| > 1, Benjamini–Hochberg adjusted *P* < 0.01, mean normalized count ≥ 5); GO enrichment used topGO (weight01, Fisher’s exact test, *P* < 0.01). Gene lists and full enrichment results are given in supplementary tables 16-18.

**Supplementary Figure 11.**
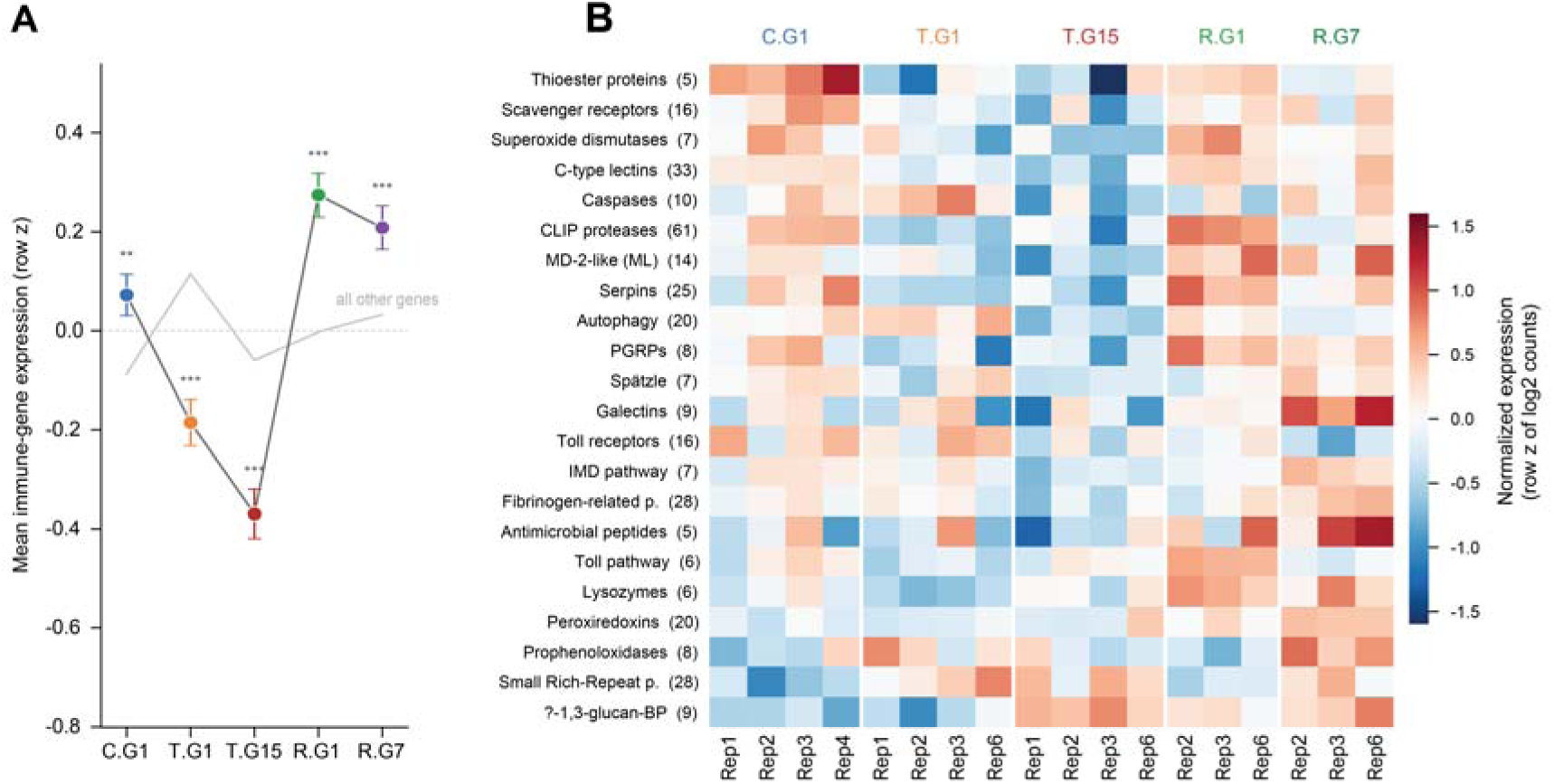
Immune gene expression is depressed by heat and restored within one generation of relaxation. (**A**) Mean normalised expression of the 360 immune genes mapped to the AalbF5 assembly, across the five sequenced conditions. Expression is the row z-score of log2 normalised counts computed across conditions on replicate-line means (the two sequencing pools per line averaged first), so zero denotes each gene’s own mean across the experiment. Coloured points show the immune-gene mean ± s.e.m.; the grey line shows all other expressed genes on the same scale. Asterisks give the significance of the deviation of the immune set from an expression-decile-matched permutation null (4,000 random gene sets of equal size): **P < 0.01, ***P < 0.001, Benjamini–Hochberg corrected. (**B**) Mean expression of each of 22 immune gene families (number of genes in parentheses) in every sequenced replicate line, grouped by condition; colour is the row z-score of log2 normalised counts computed across all samples. Families are ordered by their expression at T.G15. Most families follow the same thermal trajectory — reduced under heat, restored on relaxation — with small rich-repeat proteins and β-1,3-glucan-binding proteins as the principal exceptions. Conditions: C.G1, control generation 1; T.G1 and T.G15, tropical lines at generations 1 and 15; R.G1 and R.G7, relaxed lines at generations 1 and 7 after return to control temperature.

**Supplementary Figure 12.**
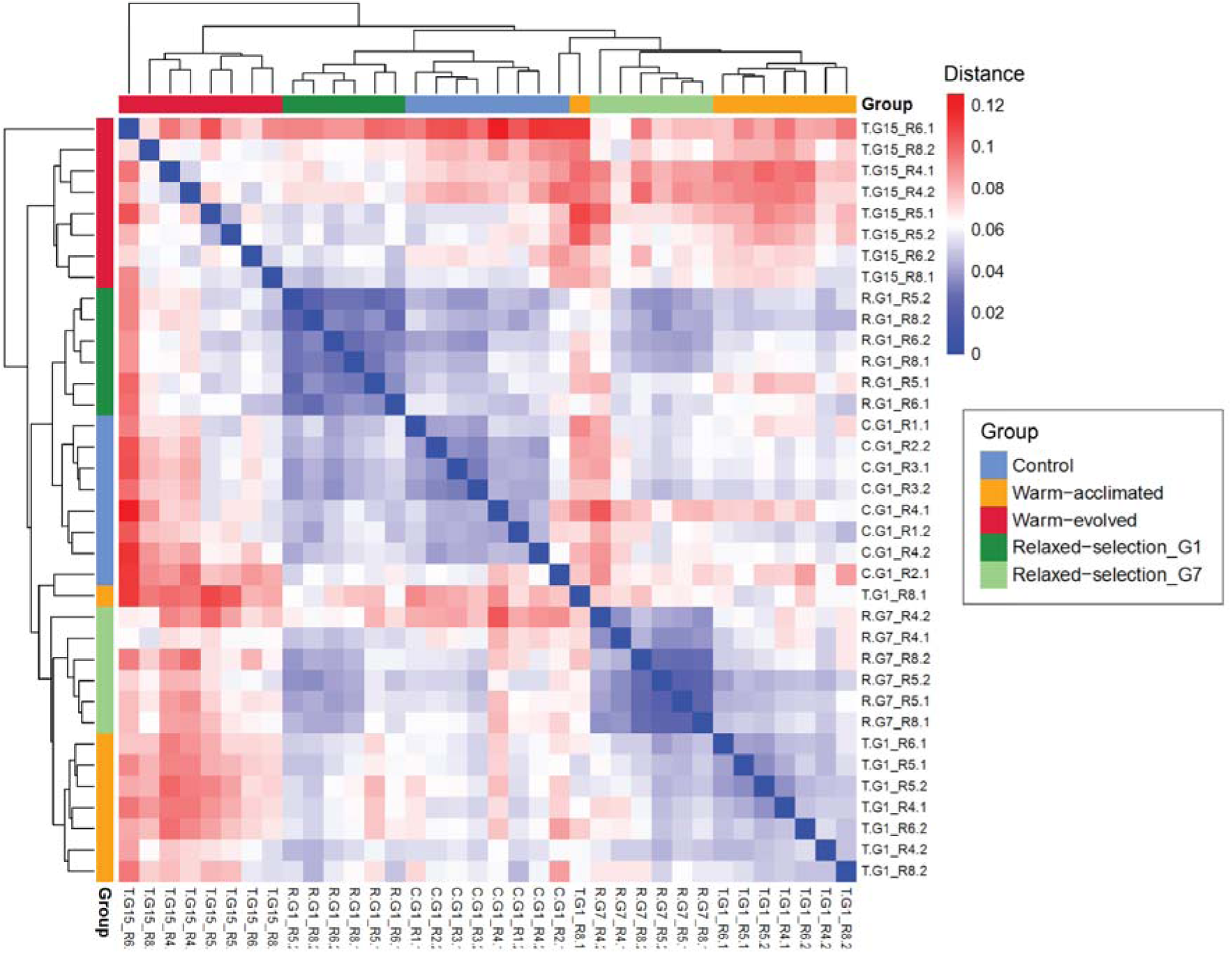
Transcriptomic similarity among experimental groups. Pairwise sample distance heatmap based on Pearson correlation distance (1 − r) of log₂-transformed DESeq2 normalized expression values. Genes with mean normalized counts below 5 were excluded prior to analysis. Rows and columns are ordered by complete-linkage hierarchical clustering (Euclidean distance); the color bar indicates experimental group: control (C.G₁, light blue), warm-acclimated (T.G₁, orange), warm-evolved (T.G₁₅, red), relaxed selection at generation 1 (R.G₁, dark green), and relaxed selection at generation 7 (R.G₇, light green). Color intensity ranges from blue (low distance, high similarity) to red (high distance, low similarity).

## Supplementary Tables Legends

Column-by-column definitions for every field of every sheet are given in the “Supplementary table summary” sheet of the data workbook; analytical parameters are given in the Materials and Methods.

**Supplementary Table 1.** Fitness traits of the control line across generations of experimental evolution. Descriptive statistics and post-hoc comparisons for 14 life-history traits measured in mosquitoes held at 28 °C at G₁, G₅, and G₁₀. Rows 4 to 36 report the traits tested by Kruskal–Wallis with Dunn’s post-hoc test (*); rows 37 onward report the traits tested by one-way ANOVA with Tukey’s post-hoc test (**). Traits include egg hatching time (EHT), larval developmental time (LDT), pupal developmental time (PDT), female and male emergence time (FET, MET), egg hatchability, larval viability, pupation rate, egg-to-adult viability, sex ratio (females), fecundity, fertility, and progeny per female.

Related to Supplementary Figure 3.

**Supplementary Table 2.** Fitness traits across thermal regimes and generations. Descriptive statistics and post-hoc comparisons for all life-history traits measured during experimental evolution across three blocks: (A) juvenile development; (B) female reproductive capacity, including wing length as a proxy for body size; and (C) adult longevity by sex and replicate line. Traits include all life-history traits detailed in Supplementary Table 1, plus wing length.

Related to Fig. 1B-1C; Supplementary Figures 1 and 2.

**Supplementary Table 3.** Energy reserves of newly emerged adults across thermal regimes and generations. Protein, glycogen, total lipid, and triglyceride content. For each metabolite, the sheet reports the omnibus one-way ANOVA, followed by group descriptive statistics and Tukey’s multiple-comparisons test results.

Related to Fig. 1D.

**Supplementary Table 4.** Thermal-behavioural traits and egg morphometry across thermal regimes and generations. Descriptive statistics and post-hoc comparisons for thermal preference, heat knockdown, egg cold tolerance, and egg shape. Traits evaluated include thermal gradient preference, knockdown temperature, egg freezing point, egg length, egg width, and egg index (length/width).

Related to Fig. 1E-F.

**Supplementary Table 5.** Genes differentially expressed during thermal acclimation (T.G₁ vs C.G₁). All 860 differentially expressed genes (DEGs) in the fat body of warm-acclimated relative to control mosquitoes. C.G₁ serves as the reference level; therefore, a positive log₂ fold change indicates upregulation in T.G₁.

Related to Fig. 2A, 2C and 2D; Supplementary Figure 4.

**Supplementary Table 6.** Co-expression modules of thermal acclimation (C.G₁ and T.G₁). WGCNA network built on control and warm-acclimated libraries (signed hybrid, soft-thresholding power 10, minimum module size 30, merge cut height 0.30). Block (A) provides module eigengene statistics, and block (B) presents the GO enrichment for each module.

Related to Supplementary Figures 4 and 5.

**Supplementary Table 7.** Recovery of fitness trait deviations from control across generations of experimental evolution. Per-trait linear slope of the standardized deviation from the pooled control (|d|) across tropical generations, for the 15 traits in the acclimation/adaptation panel, together with their assignment to one of two trajectory-shape clusters (Development / viability / longevity, or Reproductive output) from correlation-based clustering of the |d| trajectories, and each trait’s bootstrap co-clustering stability (fraction of 1,000 bootstrap resamples retaining the trait in its assigned cluster). Related to Supplementary Figure 6.

**Supplementary Table 8.** Adult mortality within one day of emergence in warm-selected lines. Percentage of adults found dead one day after emergence, categorized by generation and sex.

Related to Supplementary Figure 1.

**Supplementary Table 9.** Parameters of the daily adult-emergence curve. Best-fit parameters, along with 95% confidence intervals, of the curve fitted to the daily percentage of emerging adults, separated by thermal regime, generation, and sex.

Related to Supplementary Figure 7.

**Supplementary Table 10**. Genes differentially expressed in warm-evolved mosquitoes relative to control (T.G₁₅ vs C.G₁). All 1,387 DEGs between warm-evolved and control fat bodies; C.G₁ is the reference level. Column organization mirrors that of Supplementary Table 5.

Related to Fig. 2A and 2E.

**Supplementary Table 11**. Genes differentially expressed in warm-evolved relative to warm-acclimated mosquitoes (T.G₁₅ vs T.G₁). All 1,455 DEGs between T.G₁₅ and T.G₁ fat bodies; T.G₁ is the reference level. Column organization mirrors that of Supplementary Table 5.

Related to Fig. 2A and 2E.

**Supplementary Table 12.** Genes in each intersection of the warm-evolution UpSet plot. The 2,888 entries comprising the five intersections of the directional UpSet built on T.G₁ vs C.G₁, T.G₁₅ vs C.G₁, and T.G₁₅ vs T.G₁.

Related to Fig. 2E.

**Supplementary Table 13**. GO enrichment of the warm-evolution UpSet intersections. Enriched GO terms for each intersection presented in Supplementary Table 12, tested separately for up- and downregulated genes.

Related to Fig. 2E.

**Supplementary Table 14**. Co-expression modules of warm evolution (C.G₁, T.G₁ and T.G₁₅). WGCNA network constructed using all three groups (signed hybrid, soft-thresholding power 6, minimum module size 30, merge cut height 0.30). Block (A) details module eigengene statistics, while block (B) lists the GO enrichment for each module.

Related to Supplementary Figures 8 and 9.

**Supplementary Table 15**. Sensitivity of the R₀ trajectory to the assumed oviposition lag. Quadratic regression of R₀ on generation in the warm regime (equation 5), refitted for assumed oviposition lags (L) of 1, 2, and 3 days. The model reported in the main text utilizes L = 2.

Related to Fig. 2F.

**Supplementary Table 16**. The 83 genes differentially expressed in both thermal transitions. Genes identified as DEGs both upon the shift to the warm regime (T.G₁ vs C.G₁) and upon the return to control conditions (R.G₁ vs T.G₁₅), classified according to their expression trajectory during warm evolution. Of these 83 genes, 63 change in opposite directions in the two contrasts, and 20 change in the same direction.

Related to Supplementary Figure 10.

**Supplementary Table 17**. The 726 genes differentially expressed only on the return to control conditions (R.G₁ vs T.G₁₅). Genes that responded to the return to 28 °C but were not categorized as DEGs during the initial shift to the warm regime, comprising 363 upregulated and 363 downregulated genes.

Related to Supplementary Figure 10.

**Supplementary Table 18.** GO enrichment of the gene sets in the acclimation UpSet plot. Enriched GO terms for each gene set derived from the directional UpSet constructed on T.G₁ vs C.G₁ and R.G₁ vs T.G₁₅.

Related to Supplementary Figure 10.

**Supplementary Table 19.** Immune gene expression relative to an expression-matched permutation null, and by replicate line. Mean z-score of log2-normalized expression across the 360 fat-body-expressed immune genes at each of the five thermal conditions, tested against a null built from 4,000 expression-decile-matched random gene sets of the same size (A). Same expression values broken down by immune gene family (22 families) and individual RNA-seq replicate line, so that family-level shifts can be inspected without averaging across lines (B). Related to Supplementary Figure 11.

**Supplementary Table 20**. Reversal scores of the warm-evolution DEGs after relaxed selection. Per-gene reversal scores calculated for the 1,253 DEGs from the T.G₁₅ vs C.G₁ contrast exhibiting a |log₂ fold change| > 0.3 and a mean normalized count ≥ 10. The score, defined as 1 − LFC(R.G₇ vs C.G₁) / LFC(T.G₁₅ vs C.G₁), quantifies the extent to which expression in R.G₇ has returned toward control levels.

Related to Fig. 3B.

**Supplementary Table 21**. The 754 genes differentially expressed only after relaxed selection (R.G₇ vs C.G₁). Genes that become differentially expressed only following seven generations of relaxed selection and do not appear as DEGs in any of the warm-evolution contrasts.

Related to Fig. 3C.

**Supplementary Table 22**. GO enrichment by reversal category and for the genes specific to relaxed selection. Enriched GO terms corresponding to each reversal category defined in Supplementary Table 20, as well as for the R.G₇-only genes from Supplementary Table 21, with testing performed separately for up- and downregulated genes.

Related to Fig. 3C.

